# A comprehensive assessment of tandem repeat genotyping methods for Nanopore long-read genomes

**DOI:** 10.64898/2026.02.28.708646

**Authors:** Elbay Aliyev, Akshay Avvaru, Wouter De Coster, Garrison M. Arner, Denis M. Nyaga, Sophia B. Gibson, Ben Weisburd, Bida Gu, Claudia Gonzaga-Jauregui, 1000 Genomes Long-Read Sequencing Consortium, Mark J.P. Chaisson, Danny E. Miller, Elizabeth Ostrowski, Harriet Dashnow

**Affiliations:** Department of Biomedical Informatics, University of Colorado Anschutz, Aurora, CO, USA; Department of Biomedical Sciences, University of Antwerp, Antwerp, Belgium; VIB Center for Molecular Neurology, VIB, Antwerp, Belgium; Department of Computer Sciences, Metropolitan State University of Denver, CO, USA; Liggins Institute, University of Auckland, New Zealand; Department of Genome Sciences, University of Washington, Seattle, WA, USA; Program in Medical and Population Genetics, Broad Institute of MIT and Harvard, Cambridge, MA, USA; Department of Quantitative and Computational Biology, University of Southern California, Los Angeles, CA, USA; International Laboratory for Human Genome Research, Laboratorio Internacional de Investigación sobre el Genoma Humano, Universidad Nacional Autónoma de México, Querétaro, Mexico; Norris Comprehensive Cancer Center, University of Southern California, Los Angeles, CA, USA; Division of Medical Genetics, Department of Pediatrics, University of Washington, Seattle, WA; Department of Laboratory Medicine and Pathology, University of Washington, Seattle, WA; Brotman Baty Institute for Precision Medicine, University of Washington, Seattle, WA; Department of Biological Sciences, University of Washington Bothell, WA, USA

## Abstract

**Background:** Tandem repeats (TRs) play critical roles in human disease and phenotypic diversity but are among the most challenging classes of genomic variation to measure accurately. While it is possible to identify TR expansions using short-read sequencing, these methods are limited because they often cannot accurately determine repeat length or sequence composition. Long-read sequencing (LRS) has the potential to accurately characterize long TRs, including the identification of non-canonical motifs and complex structures. However, while there are an increasing number of genotyping methods available, no systematic effort has been undertaken to evaluate their length and sequence-level accuracy, performance across motifs from STRs to VNTRs and across allele lengths, and, critically, how usable these tools are in practice.

**Results:** We reviewed 25 available bioinformatic tools, and selected seven that are actively maintained for benchmarking using publicly available Oxford Nanopore genome sequencing data from more than 100 individuals. Our benchmarking catalog included ∼43k TR loci genome-wide, selected to represent a range of simple and challenging TR loci. As no “truth” exists for this purpose, we used four complementary strategies to assess accuracy: concordance with high-quality haplotype-resolved Human Pangenome Reference Consortium (HPRC) assemblies, Mendelian consistency in Genome in a Bottle trios, cross-tool consistency, and sensitivity in individuals with pathogenic TR expansions confirmed by molecular methods. For all comparisons, we assess both total allele length and full sequence similarity using the Levenshtein distance. We also evaluated installation, documentation, computational requirements, and output characteristics to reflect real-world use. We provide a complete analysis workflow for all tools to support community reuse.

Tool performance varied substantially across both accuracy and usability. Most methods achieved high concordance with HPRC assemblies, with higher accuracy when using the R10 ONT pore chemistry. Accuracy generally declined with increasing allele length, and most tools performed worse on homopolymers, likely reflecting underlying sequencing accuracy. Tools generally performed worse at heterozygous loci and at alleles that differed from the reference genome. Interestingly, concordance with assembly in population samples did not predict sensitivity to pathogenic expansions, with different genotypers performing best in each category. Similarly, Mendelian consistency was highest in the tool that performed worst in assembly concordance.

**Conclusions:** No single genotyper emerged as consistently best across all assessments, but strong contenders emerged in each. Our results demonstrate that length accuracy (a typical benchmarking approach) alone overestimates TR genotyping performance. Sequence-level benchmarking is essential for selecting tools best-suited for population studies and clinical diagnostics. This work provides practical guidance for tool selection and highlights key priorities for future long-read TR genotyping method development.

## Background

While tandem repeats (TRs) are among the most mutable regions in the genome, with mutation rates on the order of 10^-5^ per generation [1–4], over three orders of magnitude higher than single-nucleotide variants, they are among the most challenging classes of variation to measure accurately. TRs include short tandem repeats (STRs, 1-6 bp motifs) and variable number tandem repeats (VNTRs, 7+ bp motifs). Today, more than 70 TR loci are associated with monogenic diseases in humans [5–7], and many more are associated with increased risk of complex disease or other phenotypic variation [8–12]. TRs remain understudied due to challenges in accurately genotyping them at scale.

Genome-wide studies using short-read sequencing (SRS) approaches have improved our understanding of the scale of TR variation and allowed numerous new disease discoveries. But, while bioinformatic method development for TR genotyping using SRS has been highly active [13], these methods are limited to evaluating short, relatively simple loci due to their inability to span longer repeat alleles and fully characterize sequence variation within these alleles.

Long-read sequencing (LRS) has dramatically increased our ability to fully characterize long and complex repeat structures by revealing previously unknown features of TR biology, including motif diversity and novel mutation patterns within TR loci. This is possible because the longer read lengths generated by LRS more easily span longer repeat structures, align better within repetitive regions, and are more easily assembled than SRS data. These data can then be more easily used to resolve complex TR arrangements, assess somatic variation, phase alleles into haplotypes, and identify base modifications using a single data source. Still, challenges remain in genotyping TRs using LRS data, including higher per-base error rates, increased homopolymer errors, and, historically, sequencing cost.

Accurate determination of allele sequence, rather than just repeat number and allele size, is important for diagnosing and predicting the prognosis of many repeat expansion disorders. For example, within *RFC1*, specific motifs are considered benign at any size, while other motifs are pathogenic when expanded [14]. In the *FMR1* gene, AGG interruptions of the CGG repeat are stabilizing [14,15]. These interruptions reduce the probability that a parent with a premutation allele close to the threshold for pathogenicity will have a child with Fragile X disorder, which has important counseling implications [16,17]. The importance of uninterrupted repeats, rather than repeat length *per se*, has been shown for several of the spinocerebellar ataxias, Huntington’s Disease, myotonic dystrophy, and other repeat expansion disorders [18–24]. In these disorders, interruptions may modify the age of onset, the probability of transmission, and clinical expressivity. Collectively, these findings underscore the importance of developing accurate methods to determine not just allele lengths, but full repeat sequences with their motifs and interruptions.

There is an increasing number of long-read TR genotypers that attempt to capitalize on this rich data while taking different approaches to overcome its limitations. More than 20 long-read TR genotypers have been introduced since 2015 [25]. These tools have evolved alongside discoveries of new loci and mutation dynamics, extending beyond characterizing length to include sequence variation at the locus and methods to decompose sequences into component motifs. These tools have also improved their ability to phase variants and to model genotypes at loci with high somatic instability, while maintaining reasonable computational time and memory requirements. Despite this explosion in computational method development, little is known about the tools’ sequence-level accuracy, particularly at longer alleles and VNTRs. Most benchmarking to date has been performed by the developers of specific tools, comparing their methods to previous ones, and has often focused on allele length rather than the full allele sequence, despite sequence composition being essential for interpreting motif structure, interruptions, and pathogenic associations. Thus, there is a major gap, especially as the field increasingly relies on LRS for clinical variant discovery, interpretation, and population-scale studies. As part of the 1000 Genomes Long-Read Sequencing Consortium (1KGP-LRSC) [26], we sought to systematically evaluate and identify the most accurate algorithms to ultimately create a public reference set of TR genotypes for downstream analyses by the genomics community.

We conducted a comprehensive sequence-level benchmark of seven actively maintained long-read TR genotypers selected for their broad adoption: STRkit [27], LongTR [28], Straglr [29], STRdust [30], vamos [31], ATaRVa [32], and Medaka Tandem [33]. For the six tools that can report full allele sequences, we assessed both length and sequence accuracy. Straglr did not report sequences and was therefore assessed solely on length. For this benchmark, we used public Oxford Nanopore Technologies (ONT) whole-genome LRS data from more than 100 individuals generated by the Genome in a Bottle (GIAB) Consortium [34], the Human Pangenome Reference Consortium (HPRC) [35], and adaptive sampling data that targeted relevant genes of individuals with pathogenic expansions [36]. We genotyped ∼43,000 TR loci from the TRExplorer catalog [37], selected to span a wide range of motif lengths, repeat lengths, repeat complexity, and genomic contexts. Because no genome-wide ground truth exists, we benchmarked accuracy using four complementary strategies: concordance with haplotype-resolved, high-quality HPRC assemblies; Mendelian consistency in a GIAB trio; cross-tool agreement; and sensitivity to pathogenic alleles confirmed by repeat-primed PCR (RP-PCR) and Southern blot. We evaluated both allele length and full-sequence similarity using the Levenshtein distance and assessed installation, compute requirements, documentation completeness, and output formats to reflect real-world usability. We discuss their strengths not only in terms of accuracy and performance, but also in the information they provide for each locus to support downstream analysis, and critically, their usability, from installation to execution and output parsing. Finally, we discuss current limitations and major pain points for users, highlighting areas for future development.

## Results

### Genotyper selection

We reviewed the literature and Long Read Tools resource [38,39] to identify 25 long-read TR genotyping algorithms (**Supplementary Table 1**). From these, we selected open-source tools that had at least one update since 11 November 2025 that accepted ONT sequencing reads and genotyped both STRs and VNTRs. We further narrowed our selection to genotypers that can output full allele sequences in Variant Call Format (VCF), allowing users to identify motif sequences, interruptions, and compound allele structures. For comparison, we included Straglr, the most popular length-only genotyper. The final selection included eight tools (**Supplementary Table 2**), and of these, we were able to install and run seven (see Usability): STRkit [27], LongTR [28], Straglr [29], STRdust [30], vamos [31], ATaRVa [32], and Medaka Tandem [33]. We were unable to install TRsv [40] without substantial code modification, so we removed it from the study. Note that TRGT [41] was excluded as it does not accept ONT data.

**Table 1.**
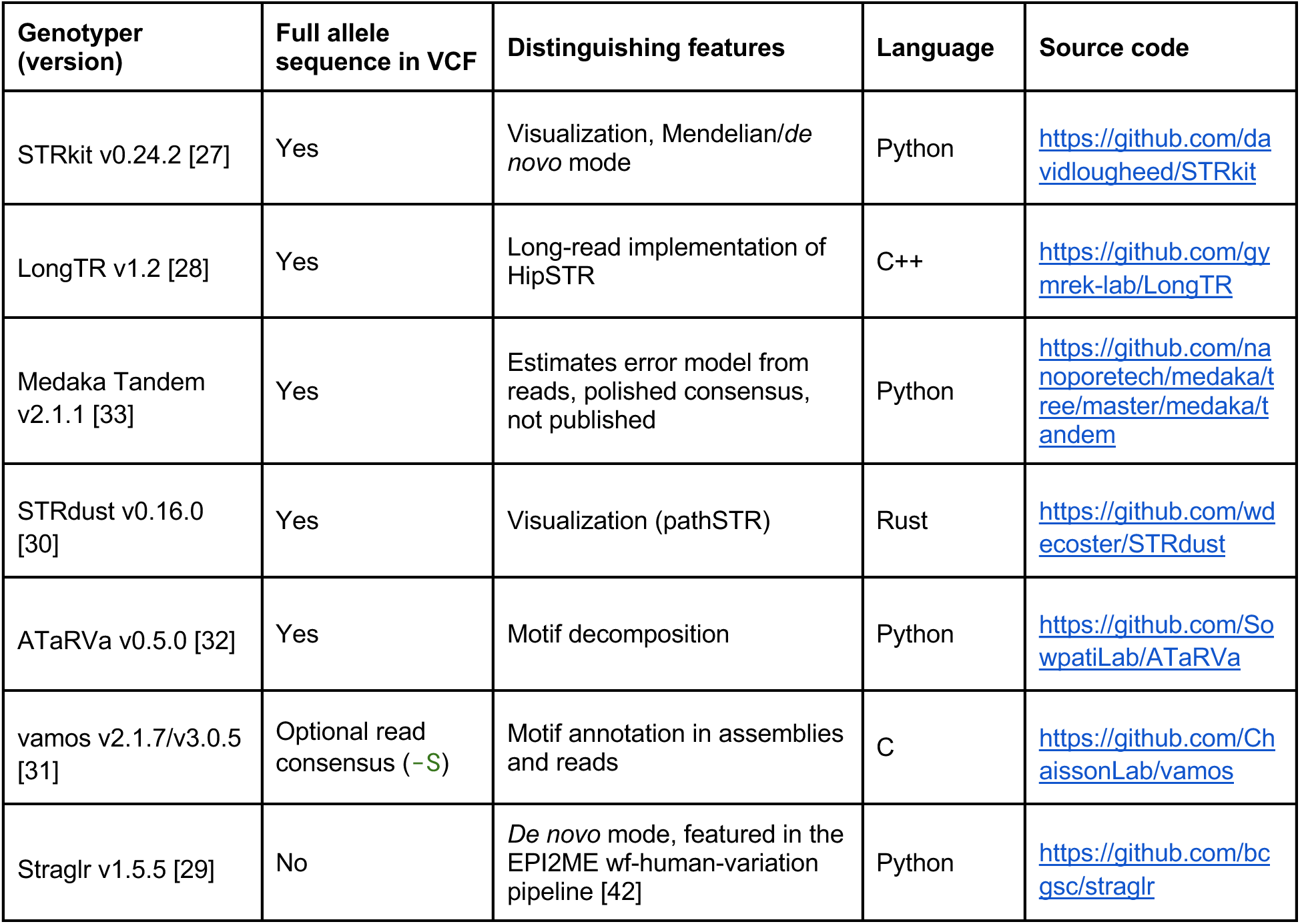
Tools evaluated in this study with key distinguishing features.

### Usability

#### Catalog generation

Unlike most modern SNV, short indel, and SV genotyping methods, TR genotyping is typically done at a pre-defined set of loci based on an annotation of the reference genome. The genotypers we assessed all require such a catalog, often in Browser Extensible Data (BED) format and with slightly varying formats (see **Methods**). Most use a 0-based standard first three BED columns: chromosome, start position, end position, followed by a fourth motif sequence column. LongTR uses 1-based start coordinates in the catalog, while vamos uses a 1-based coordinate system by default, but can accept 0-based with the -Z flag. Tools typically provide their own appropriately formatted catalog for GRCh38, often including known pathogenic loci and/or broader genomic catalogs based on TRF annotations. To assess tools using a common catalog, we developed a benchmark catalog of 43,009 loci by sampling isolated loci from the TRExplorer catalog [37] (see **Methods**). As the human genome is disproportionately rich in smaller motifs, and to fully explore this metric, we designed the catalog to sample up to 1000 loci for each motif size, then supplemented it with the 75 disease-associated loci in STRchive [5].

#### Genotyper installation and user experience

All benchmarking workflows were implemented using the Nextflow workflow management system (DSL2), enabling scalable and reproducible execution across high-performance computing environments. Each Nextflow process was defined within a fully self-contained environment, avoiding reliance on inherited shell modules and preventing clashes between dependency versions, ensuring consistent behavior across compute nodes. Tool dependencies were managed using either Singularity container images or Conda environments, depending on software compatibility: LongTR, Straglr, and STRkit were deployed via containers (Wave/Seqera community images or GitHub Container Registry), while ATaRVa, Medaka, and vamos required Conda environments; STRdust was run as a statically pre-compiled binary. Several software bugs and compatibility issues were encountered and reported during the course of this work. For Medaka, we identified two bugs: first, when input CRAM files contained a UR tag in the BAM header pointing to a reference path that no longer existed on the local filesystem, Medaka ignored the user-supplied reference and attempted to use the path embedded in the header, causing the run to fail ([issue #574](https://github.com/nanoporetech/medaka/issues/574#issuecomment-3841481587)); second, when Medaka tandem was run on certain GIAB samples (HG002 and HG006) it would process ∼98% of loci and then hang indefinitely without completing or terminating, exhausting wall-time limits and leaving large numbers of temporary files behind ([issue #583](https://github.com/nanoporetech/medaka/issues/583#issuecomment-3946647710)). For vamos, the TRExplorer catalog could not be used directly because vamos does not support overlapping loci and raises a hard error when the start coordinate of a locus precedes the end coordinate of the previous one; critically, even the vamos-specific catalog exported from TRExplorer still contained such overlapping entries, which required the TRExplorer team to issue a corrected catalog before it could be run ([issue #10](https://github.com/broadinstitute/TRExplorer/issues/10#issuecomment-3629331647)).

#### VCF output formats

The usability of a tool depends largely on ease of interpretation and integration with downstream analyses. The VCF format, originally developed for reporting simple genomic variants, was extended in version 4.4 to include TR specifications [43]. However, this extension has not been widely adopted. Before this update, TR variant callers for SRS platforms relied on flexible adaptation of the standard VCF format, and these conventions were largely carried forward by long-read genotypers. As a result, output formats vary across tools, sometimes dramatically.

Most tools report VCF results, although some also provide auxiliary output files. Straglr generates a tab-separated values (TSV) file containing read-level information for each locus and a summary variants file in BED format. STRkit outputs an overview in TSV format, along with a more extensive JavaScript Object Notation (JSON) report. Medaka Tandem produces multiple supplementary files, including reports of missing regions, consensus allele sequences, and binary alignment map (BAM) files of reads mapped to consensus alleles.

LongTR, ATaRVa, STRkit, Medaka Tandem, and STRdust all produce VCF output with complete allele sequences in REF and ALT fields, while using distinct keys in FORMAT and INFO fields. Straglr instead reports allele lengths in the INFO field. vamos includes most relevant information within the VCF with the addition of the -S parameter, but its formatting differs from that of other tools and generally requires additional reformatting for compatibility (see **Methods**).

#### Computational Resources

Computational efficiency is becoming increasingly critical as analyses move to population scales in cloud compute environments, where even modest differences in resource requirements can be costly. We developed a Nextflow [44] pipeline to run all genotypers and record computational resource usage (https://github.com/dashnowlab/TR-Benchmarking). Tools were run on one of the University of Colorado Boulder Alpine’s [45] x86_64 AMD Milan CPU in single-thread mode with 32GB of physical memory, which was increased to 64GB for failed runs (Straglr only) **(Supplementary Table 3)**. We attempted to run all genotypers on the ∼5 million loci in the full TRExplorer Catalog; however, Alpine limits jobs to one week, and some tools were still running after seven days. We therefore focused our subsequent benchmarking efforts on the described curated 44k catalog. Physical memory usage was modest for most tools: 1-1.5 GB per sample for vamos 3.0.5, ATaRVa, and STRdust. STRkit, LongTR, and Medaka Tandem typically required 2.5-3.5 GB, with LongTR using up to 6 GB for some samples. Straglr had a much higher mean memory usage of 37.3 GB, with up to 70 GB for one sample. Vamos was the fastest (mean 73 mins), closely followed by STRkit (90 mins) and LongTR (98 mins). Runtimes for ATaRVa, Medaka Tandem, and STRdust were typically around 200 minutes. Straglr’s mean runtime was 13.5 hours per sample, with a maximum of almost 20 hours. All tools can be run with multi-threading to substantially reduce elapsed runtime, except for LongTR, which can be run by chromosome or other subsets, followed by merging, but this must be handled by the user.

### Accuracy

For TR genotyping, ground truth is elusive as there is no single gold-standard reference method that can fully resolve both length and sequence composition at scale. Therefore, we devised four strategies to assess accuracy: assembly concordance, Mendelian consistency, genotyper consistency, and sensitivity to detect pathogenic expansions confirmed by orthogonal molecular techniques (**Figure 2**). Two comparison metrics were used: 1) we compared total allele length in bp, and 2) assessed full sequence similarity using Levenshtein distance to calculate the number of edits between two alleles.

**Figure 1.**
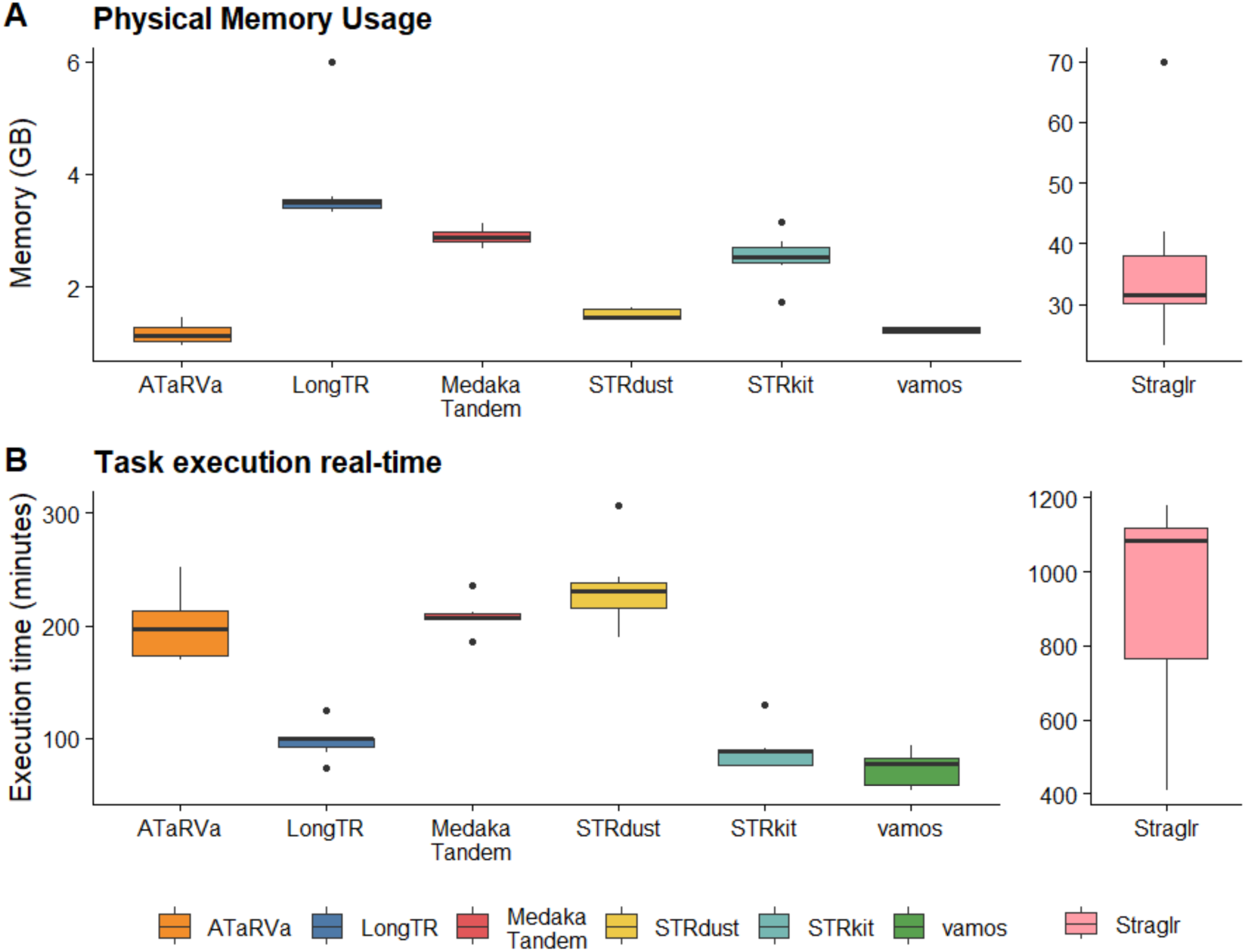
The seven GIAB samples were subsampled to ∼30X and genotyped at 43,009 TR loci. Tools were run on one of CU Boulder Alpine’s x86_64 AMD Milan CPU in single-thread mode with 32GB of physical memory. STRaglr failed at 32 GB of memory and re-run with 64GB of memory. **A.** Memory describes the peak physical RAM (peak resident set size) per sample, while **B.** Execution time is the elapsed run time for the genotyping task on a single thread.

**Figure 2.**
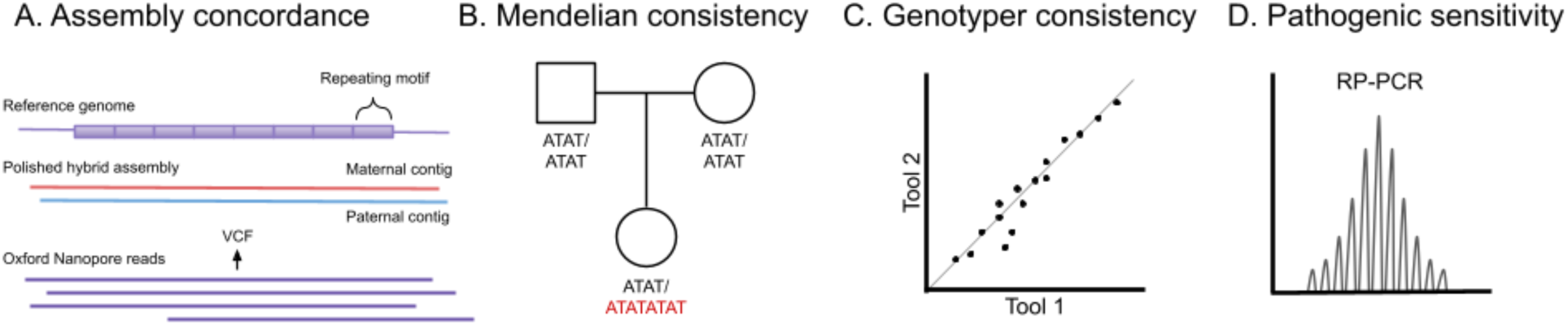
Genotyping tools were run on 30-35x ONT reads aligned to GRCh38 to assess assembly concordance, Mendelian consistency, genotyper consistency, and sensitivity to pathogenic expansions. Two comparison metrics were used: 1) length, and 2) full sequence using Levenshtein distance. **A**. Polished hybrid ONT/HiFi assemblies from HPRC were aligned to **GRCh38** to assess assembly concordance. For each tool’s VCF record, we extracted the corresponding alleles from the assembly. **B.** In the two GIAB trios, probands were compared to parents at each locus to assess Mendelian consistency. Any Mendelian inconsistency (i.e., apparent *de novo* mutation) was assumed to be an error. **C.** For each pair of tools, genotyper consistency was assessed using allele similarity at loci called by both tools. **D.** Tools were run on data collected using adaptive sampling on the ONT platform from samples with known pathogenic alleles confirmed by repeat-primed PCR (RP-PCR) and, in some cases, Southern blot. Results were then assessed for sensitivity based on their ability to identify an allele in the disease pathogenic range.

#### Assembly concordance

Genome assembly provides an orthogonal approach to read-based genotyping. It is still unclear if assembly- or read-based methods are more accurate at resolving tandem repeat loci; however, assemblies constructed from deep coverage combining multiple long- and short-read sequencing technologies are expected to yield greater accuracy than average ONT alignments alone. We used polished hybrid ONT/HiFi assemblies generated by the HPRC, which integrated 60x PacBio HiFi data with 30x ultra-long (>100 kbp) ONT data and phasing using Illumina trio-WGS or Hi-C [46]. To benchmark read-based genotyping against assemblies, we downloaded ∼30-35x ONT reads from HPRC for 67 samples: 33 samples with data generated using the R9.4.1 chemistry and 34 using the R10.4.1 chemistry, along with their corresponding hybrid assemblies. We aligned the assemblies and ONT data to the GRCh38 human reference genome. For each tool’s VCF record, we extracted the corresponding alleles from the sample assembly using a liftover procedure (see Methods). Note that some tools (STRkit, LongTR) extend the coordinates from those in the catalog; therefore, we chose not to simply extract the catalog coordinate from the assembly and use a method such as Truvari [47].

For each genotype, alleles called by each tool were paired with assembly alleles such that the combined deviation or error was minimized. For each allele, the absolute deviation from the corresponding assembly allele length was calculated and categorized into four classes: *Match* (no error), *Off by 1bp* (error of exactly 1 bp), *Off by motif* (error less than or equal to the motif length of the locus), and *Mismatch* (error exceeded the motif length). LongTR applies a default minimum mean base Phred score threshold of 30 per read and had no calls in samples sequenced with R9.4.1 chemistry, as the mean read base quality for these samples falls below this threshold. The accuracy results are thus presented separately for the R10.4.1 and R9.4.1 samples.

While all tools were provided with the same ∼43k catalog, most genotypers called a mean of ∼42k loci per sample except Straglr (R10: 36,966 loci; R9: 38,019 loci) and STRkit (R10: 29,788 loci; R9: 30,011 loci) (**Supplementary Table 4**). The missing calls in STRkit could be attributed to a normalized map quality threshold of 0.9, which filters out many supporting reads.

We calculate accuracy for a tool as the percentage of called loci that fall into the deviation classes Match and Off by 1 bp in a sample. The mean accuracy of the tools, ranked from best to worst by R10 accuracy was: LongTR (R10: 98.67%), Medaka Tandem (R10: 98.61%; R9: 97.77%), ATaRVa (R10: 98.48%; R9: 97.35%), vamos (R10: 98.09%; R9: 95.94%), and STRkit (R10: 97.72%; R9: 96.21%) **(Figure 3A)**. Beyond these five tools, there is a notable drop in accuracy with Straglr (R10: 90.09%; R9: 88.22%) and STRdust (R10: 72.56%; R9: 63.56%) **(Figure 3A)**. However, when an off-by-motif deviation is permitted, the accuracy of STRdust increases to 94.93% (**Supplementary Table 5**).

**Figure 3:**
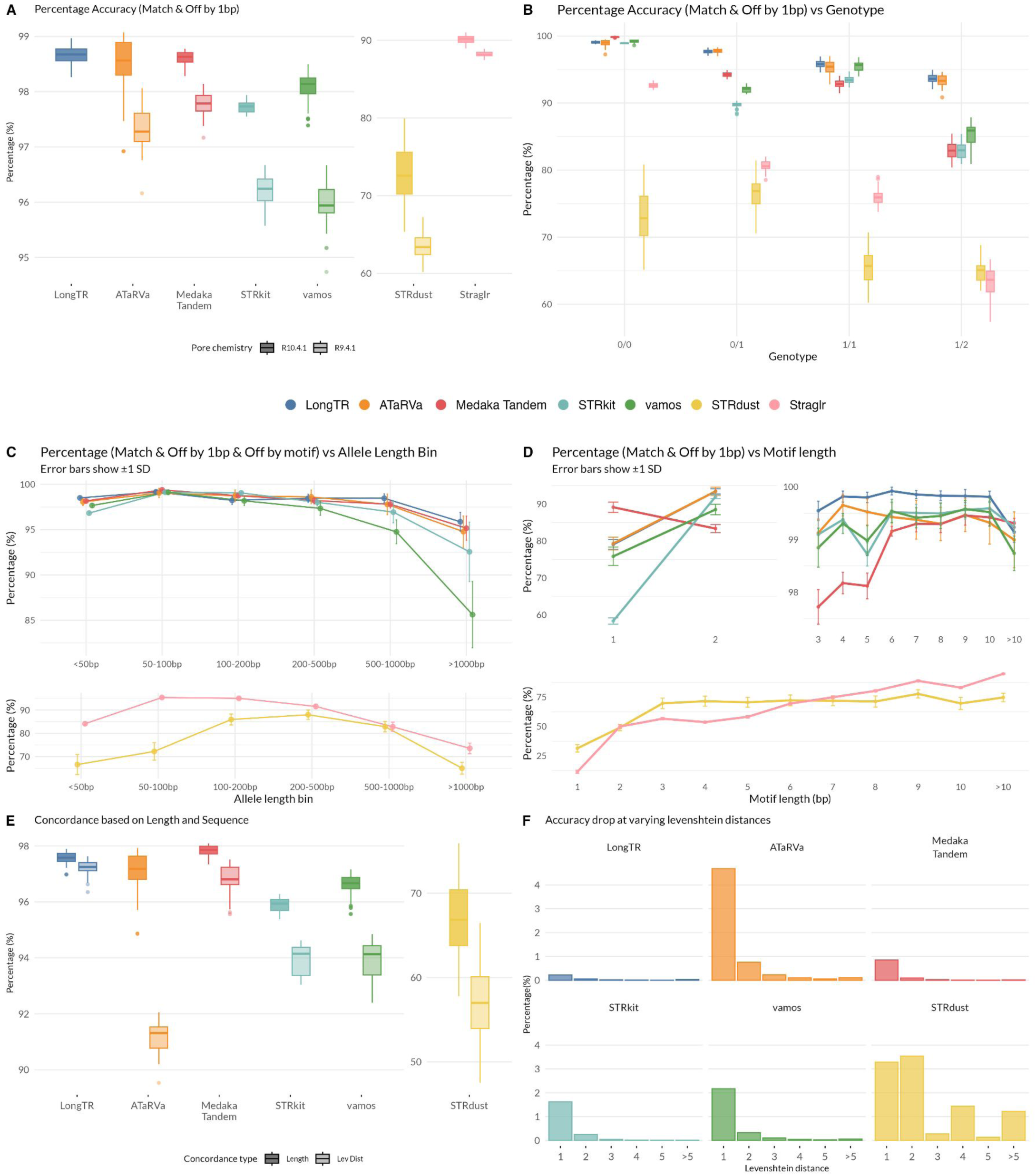
Accuracy of genotypers across different stratifications calculated as the percentage of allele calls with length within 1 bp with the assembly. **A.** Box plots show the distribution of accuracy for each tool, stratified by pore chemistry (R10.4.1, solid boxes; R9.4.1, hatched boxes). Tools are split across two y-axes for readability. **B.** Accuracy of each tool stratified by genotype class: homozygous reference (0/0), heterozygous reference (0/1), homozygous alternate (1/1), and heterozygous alternate (1/2). Points represent mean accuracy per genotype class across all samples. Points represent mean accuracy, and error bars indicate ±1 standard deviation across samples. **C.** Accuracy of each tool across allele length bins (<50 bp, 50–100 bp, 100–200 bp, 200–500 bp, 500–1000 bp, and >1000 bp) in R10.4.1 samples. Points represent mean accuracy, and error bars indicate ±1 standard deviation across samples. Tools are displayed in two separate panels for readability. **D.** Accuracy of each tool across motif length categories (1–10 bp and >10 bp) in R10.4.1 samples. Points represent mean accuracy, and error bars indicate ±1 standard deviation across samples. Tools are displayed across three panels for readability. The upper two panels show the five higher-performing tools (LongTR, ATaRVa, Medaka Tandem, STRkit, and vamos), with mononucleotide and dinucleotide repeats shown separately on a y-axis range of 50–95% and motif lengths ≥3 bp displayed on an expanded y-axis of 98–100%. The lower panel shows STRdust and Straglr on a broader scale to capture their wider accuracy range across all motif lengths. E. Accuracy calculated for each tool on two levels: length-based accuracy, considering all calls with a matching allele length, and sequence-based accuracy, calculated using Levenshtein distance for the subset of calls with a matching allele length. The difference between the two values reflects the proportion of length-concordant calls that differ at the sequence level from the assembly allele. **F.** Contribution of Levenshtein distance categories to percentage-point sequence-level accuracy drop. Bar plots showing the percentage of total calls at each Levenshtein distance value for each tool, restricted to calls with matching allele lengths. Each bar represents the proportion of calls at a given Levenshtein distance, illustrating the relative contribution of different magnitudes of sequence deviation to the overall sequence-level accuracy drop observed for each tool.

On average, 83.6% of loci in the R10 HPRC assemblies were invariant (homozygous for the reference allele). To assess tool performance at variable loci, we excluded loci with homozygous reference calls in assemblies. STRdust exhibited the smallest percentage-point drop in accuracy between all loci and non-reference loci at just 1.42%; however, this should be interpreted in the context of its lower overall accuracy relative to other tools. LongTR and ATaRVa showed near-identical drops of 2.15% and 2.17%, respectively, followed by progressively larger drops in vamos (5.91%), Medaka Tandem (6.43%), STRkit (7.69%), and Straglr (13.47%) **(Supplementary Figure 1)**.

To further characterize tool performance, we categorized variant loci by genotype: heterozygous reference (0/1), homozygous alternate (1/1), and heterozygous alternate (1/2), and assessed each tool’s performance within these groups. In the assemblies, the average proportions of genotype calls are 83.70% (0/0), 8.71% (0/1), 5.21% (1/1), and 2.36% (1/2). Most tools show broadly similar genotype proportions to the assemblies, with a few exceptions **(Supplementary Figure 2)**. STRdust tends to call more genotypes with an alternate allele, while ATaRVa and Straglr show slightly elevated 0/0 call rates (88.81% and 88.21%, respectively). STRkit, Medaka Tandem, and Straglr have fewer heterozygous (0/1 and 1/2) calls than assembly, with correspondingly higher 1/1 calls, suggesting a slight bias toward homozygous calls.

Across all tools, accuracy consistently decreases with increasing numbers of non-reference alleles **(Figure 3B)**. STRkit and vamos are exceptions, showing higher accuracy for 1/1 than 0/1 genotypes, potentially reflecting a homozygous calling bias. LongTR and ATaRVa are the most robust, maintaining accuracy above 93% even for 1/2 calls. For 0/0 genotypes, all tools except STRdust achieved >90% accuracy, with five tools approaching 98%. The per-genotype ranking broadly mirrors the overall accuracy order of LongTR, ATaRVa, Medaka Tandem, vamos, STRkit, Straglr, and STRdust, with two notable exceptions: Medaka Tandem was the top performer for 0/0 genotypes (99.83%), and ATaRVa for 0/1 genotypes (97.76%).

We next assessed tool accuracy as a function of total allele length, binning alleles into six categories: <50 bp, 50–100 bp, 100–200 bp, 200–500 bp, 500–1000 bp, and >1000 bp. Accuracy generally decreased with increasing allele length for loci above 100 bp, consistent with the expectation that longer alleles are likely to accumulate more sequencing errors **(Figure 3C)**. All tools also showed poorer accuracy for loci below 50 bp compared to the 50-100bp bin, likely because this bin contains a disproportionate number of homopolymers. Examining the number of calls per category **(Supplementary Figure 3)**, STRkit had the fewest calls across all length categories, with a proportionally increasing rate of missing calls at greater lengths. In contrast, despite having the second-highest overall rate of missing calls, Straglr concentrated most of its missing calls in the shortest length category and had the highest number of calls for alleles >1000 bp. All tools produced call counts comparable to the assemblies across most length categories, with the notable exception of the >1000 bp category, which tools tended to under-call. Among tools, Straglr, STRdust, and vamos had the highest number of calls in this longest category. When considering the absolute count with accurate calls across all tools and length categories, performance was largely comparable, with Straglr showing a slight advantage in the >1000 bp category **(Supplementary Figure 4)**.

While above we considered total allele size, we also evaluated each tool’s ability to detect expansions relative to the reference across a range of size categories: <50 bp, 50–100 bp, 100–200 bp, 200–500 bp, and >500 bp **(Supplementary Figure 5)**. All tools showed a gradual decline in accuracy with increasing expansion length, with a more pronounced drop for expansions >1000 bp. A consistent dip in accuracy was also observed across all tools in the 100–200 bp bin. ATaRVa and LongTR were the top two tools overall, with ATaRVa performing the best up to 500 bp expansions and LongTR performing better beyond that threshold. For the largest expansion category (>1000 bp), LongTR achieved the highest accuracy at 89.66% while STRkit had the lowest at 24.37%.

We next evaluated tool performance across motif length categories. Recall that the catalog was constructed by sampling 1,000 loci per motif length. The total number of calls per category was broadly consistent across tools, except for Straglr and STRkit. Straglr showed a proportionally uniform distribution of missing calls across motif length categories, whereas STRkit showed a progressively increasing rate of missing calls with longer motifs **(Supplementary Figure 6)**.

When comparing accuracy across tools, a clear split emerges: STRdust and Straglr perform markedly worse than the remaining five tools across motif lengths **(Figure 3D)**. Mononucleotide (homopolymer) and dinucleotide repeats posed the greatest challenge across all tools, likely reflecting high read-level error or possibly mosacism at these repeat types. Notably, Medaka Tandem achieved the highest accuracy for homopolymers at 89.13%, nearly 10% better than the next best tools, ATaRVa (79.31%) and LongTR (79.07%), and Straglr showed the poorest accuracy at 11.34%. **(Figure 3D)**. However, Medaka Tandem dropped to the lowest accuracy among the top five tools for dinucleotide repeats (83.25%), while the other four tools performed better (88–93.5%), with LongTR and ATaRVa performing the best. From motif length 3 bp onward, all five tools converge to accuracies of 98% or above, remaining largely stable through longer motif lengths before a slight decline in the >10 bp category. Both STRdust and Straglr improve with increasing motif length, though their trajectories differ (**Figure 3D**).

Beyond length-based accuracy, we assessed each tool’s sequence-level accuracy by comparing tool allele sequences against the assembly using Levenshtein distance. Straglr was excluded from this analysis as it does not report allele sequences. To isolate sequence-level deviations, we restricted the analysis to loci where the tool length and assembly length matched, then compared the proportion of calls with matching sequences (Levenshtein distance of 0, **Figure 3E)**. LongTR demonstrated the greatest sequence-level robustness, achieving 97.20% sequence accuracy with a drop of only 0.36% from its length-based accuracy. Medaka Tandem was the next-best performer, with a drop of 1.01%, followed closely by STRkit (1.95%) and vamos (2.72%). ATaRVa and STRdust showed notably larger deviations, with drops of 5.92% and 9.89%, respectively. Examining the distribution of Levenshtein distances at length-concordant loci, the accuracy drop for most tools is largely explained by calls with a Levenshtein distance from the assembly of 1 (i.e., a single bp SNV, **Figure 3F**). STRdust is a notable exception as it generated calls with larger Levenshtein distances, suggesting a higher prevalence of more substantial sequence-level errors.

When analyses were repeated using R9.4.1 data (except for LongTR), all tools experienced a drop in performance, though the overall ranking trends were largely preserved across metrics. ATaRVa and Medaka Tandem competed closely for the top positions, with each taking the lead in different analyses, while vamos and STRkit similarly traded places in different analyses, followed by Straglr and STRdust (**Supplementary Fig. 7**). One notable exception was STRkit, which outperformed all other tools for alleles >1000 bp in R9 data (81.83%; **Supplementary Fig. 7B)**, in contrast to its fourth-place ranking in the R10 analysis.

#### Mendelian consistency

To assess the accuracy of tandem repeat genotypes produced by each tool, we evaluated Mendelian consistency in a parent-offspring trio, reasoning that concordance with expected inheritance patterns provides an orthogonal measure of genotyping fidelity independent of the reference genome. In the HG002 trio, the proband was compared to parents at each genotyped locus, and any Mendelian inconsistency (i.e., apparent *de novo* mutation) was assumed to be an error. Because the *de novo* mutation (DNM) rate for TRs is estimated to be on the order of 10^-5^ per locus per generation [1–4], we would expect fewer than 1 DNM per generation from our catalog of 43k loci, meaning that the vast majority of apparent DNMs are likely to be errors.

We evaluated Mendelian consistency in the GIAB HG002 trio by assessing whether the child’s genotype could be explained by parental alleles, allowing for exact allele matches, calls differing by one base pair, or calls differing by up to one full repeat motif unit; a composite metric that accounts for the rounding and boundary ambiguities inherent to TR genotyping. Loci were stratified by the child’s genotype class (0/0, 0/1, 1/1, and 1/2) and by repeat motif length.

The total number of genotyped loci varied substantially across tools, which is a critical consideration when interpreting consistency metrics. Several tools genotyped similar numbers of loci (∼41.5k; Medaka Tandem, ATaRVa, LongTR, and STRdust), with vamos (n=40,173) and STRkit (n=39,131) calling slightly fewer, and Straglr (n=33,434) notably fewer. The difference is particularly pronounced at compound heterozygous loci (1/2), where STRdust called the most (n=1,955), followed by LongTR (n=1,147), STRkit (n=544), vamos (n=538), ATaRVa (n=558), and Medaka Tandem (n=692), while Straglr called only 128. This means Straglr’s consistency estimates at 1/2 loci are based on a much smaller and potentially less representative set, and its apparent accuracy may partly reflect more selective genotyping (Figure 4A).

**Figure 4.**
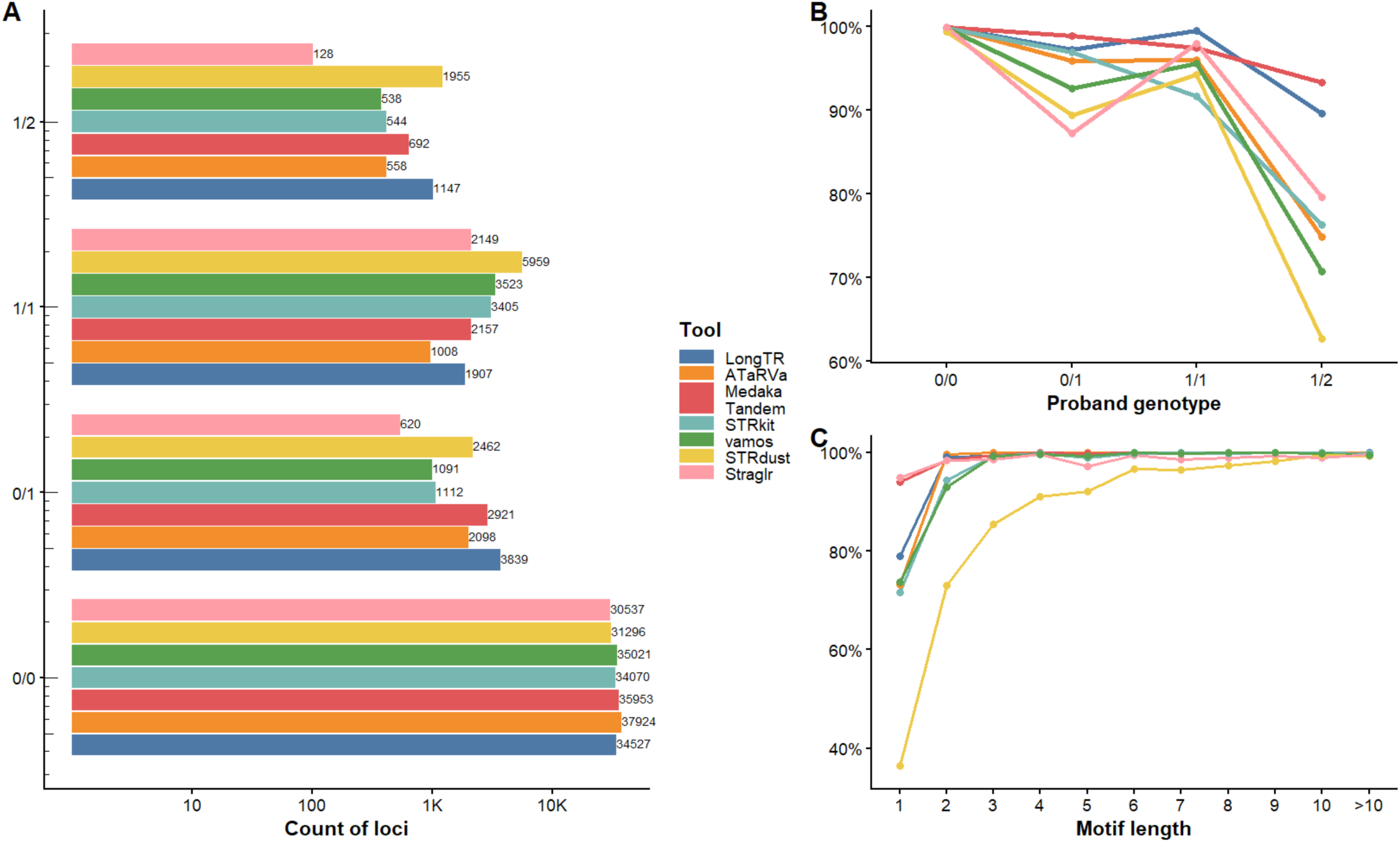
Mendelian consistency and motif-length performance across STR genotyping tools in the HG002 trio. **A)** Number of genotyped loci across four child genotype classes (0/0, 0/1, 1/1, and 1/2) for each tool. Counts are shown on a log10 scale. Values next to bars indicate the total number of successfully genotyped loci per genotype and tool. **B)** Mendelian-consistency rate (percentage of loci where the proband alleles are within one motif length of the parents) stratified by child genotype. **C)** Mendelian-consistent rate as a function of STR motif length. Motif lengths greater than 10 bp are collapsed into a single bin.

At homozygous reference loci (0/0), nearly all tools achieved >99.8% consistency (ATaRVa, Straglr: 99.99%; LongTR: 99.95%; Medaka Tandem: 99.93%; STRkit: 99.81%; vamos: 99.86%), indicating that false positive heterozygous calls are rare across methods. STRdust was the notable exception at 99.5%, driven by a higher rate of motif-length-off calls at this class (**Supplementary Table 6, Supplementary Figure 8**). Consistency declined as genotype complexity increased, with the sharpest divergence among tools observed at compound heterozygous loci (1/2), which represent the most challenging class. Here, Medaka Tandem remained the top performer at 93.4%, followed by LongTR (89.6%), Straglr (79.7%), STRkit (76.3%), ATaRVa (74.9%), and vamos (70.8%), while STRdust showed the lowest consistency at 62.7% (**Figure 4B**). At heterozygous loci (0/1), Medaka Tandem again led (98.9%), followed by LongTR (97.3%), STRkit (96.9%), ATaRVa (96.0%), vamos (92.6%), STRdust (89.4%), and Straglr (87.3%). At homozygous alternate loci (1/1), LongTR achieved the highest consistency (99.5%), followed by Straglr (97.9%), Medaka Tandem (97.5%), ATaRVa (96.0%), vamos (95.6%), STRdust (94.3%), and STRkit (91.7%).

Mendelian consistency also varied by repeat motif length (**Figure 4C, Supplementary Table 7**). Mononucleotide repeats were the most difficult to accurately call, with STRdust below 40% consistency, and ATaRVa, STRkit, and vamos ranging from 71% to 74%, reflecting the well-known higher homopolymer error in LRS. Medaka Tandem (94.1%) and Straglr (94.9%) performed notably better at mononucleotide loci. Performance improved sharply for di- and trinucleotide repeats and largely plateaued at ≥4 bp motifs, where most tools converged on 95-100% consistency. STRdust showed the steepest improvement trajectory across motif lengths, rising from below 40% at mononucleotides to above 99% at longer motifs, yet its overall consistency remains constrained by its weaker performance on shorter motifs and heterozygous genotypes.

Overall, Medaka Tandem demonstrated the most robust Mendelian consistency across all genotype classes and motif lengths in the HG002 trio, with LongTR as a strong second. Straglr performed competitively, particularly at 1/2 and 1/1 loci, though its smaller genotyped set limits direct comparison. ATaRVa, STRkit, and vamos formed a competitive intermediate tier at most genotype classes, while STRdust showed the most tool-specific weaknesses, particularly at heterozygous and compound heterozygous loci and short motifs despite genotyping the broadest set of loci.

#### Genotyper consistency

Consistency across different genotyping methods does not necessarily reflect accuracy; however, such comparisons are useful to the community for selecting a genotyping method and for assessing the usefulness of a control cohort genotyped with a different method. Consistency across callers can also point to specific loci that are potentially problematic, owing to high levels of disagreement, and may help diagnose the causes of error. Additionally, when running multiple genotypers to improve sensitivity, consistency analysis may inform the selection of complementary tools.

For each pair of tools, we assessed the similarity of allele sequences and lengths at the set of loci called by both tools. For comparisons with STRaglr, only length was considered. Genome assemblies are treated as an additional genotyping method, with genotype calls derived from the assembly based on the original catalogue coordinates. For each pair of tools, we calculated the Levenshtein distance and the absolute length difference between alleles across all commonly called loci within a sample, and averaged these values across all loci and samples. The pairwise average values, including comparisons with assembly calls, are visualized as a heatmap **(Figure 5)**, where the upper triangle shows the average Levenshtein distance, and the lower triangle represents average absolute length difference. Examining the average length difference first, LongTR, ATaRVa, Medaka Tandem, and vamos show high similarity with average length differences below 1. The increased average length difference observed between STRkit and the assembly, despite STRkit’s high overall assembly concordance, is attributable to STRkit adjusting locus coordinates, while the comparisons here are done based on original catalogue coordinates. Consistent with the overall assembly concordance results, Straglr shows the second-lowest deviation from the assembly, while STRdust shows the highest. When the assembly is included as a comparator, the average distance of each tool from the assembly is broadly comparable to, and in most cases slightly lower than, pairwise tool-to-tool distances. Notably, ATaRVa shows the smallest average length difference from the assembly, despite ranking third in overall assembly concordance. The average Levenshtein distance shows trends largely similar to those of the average length difference.

**Figure 5.**
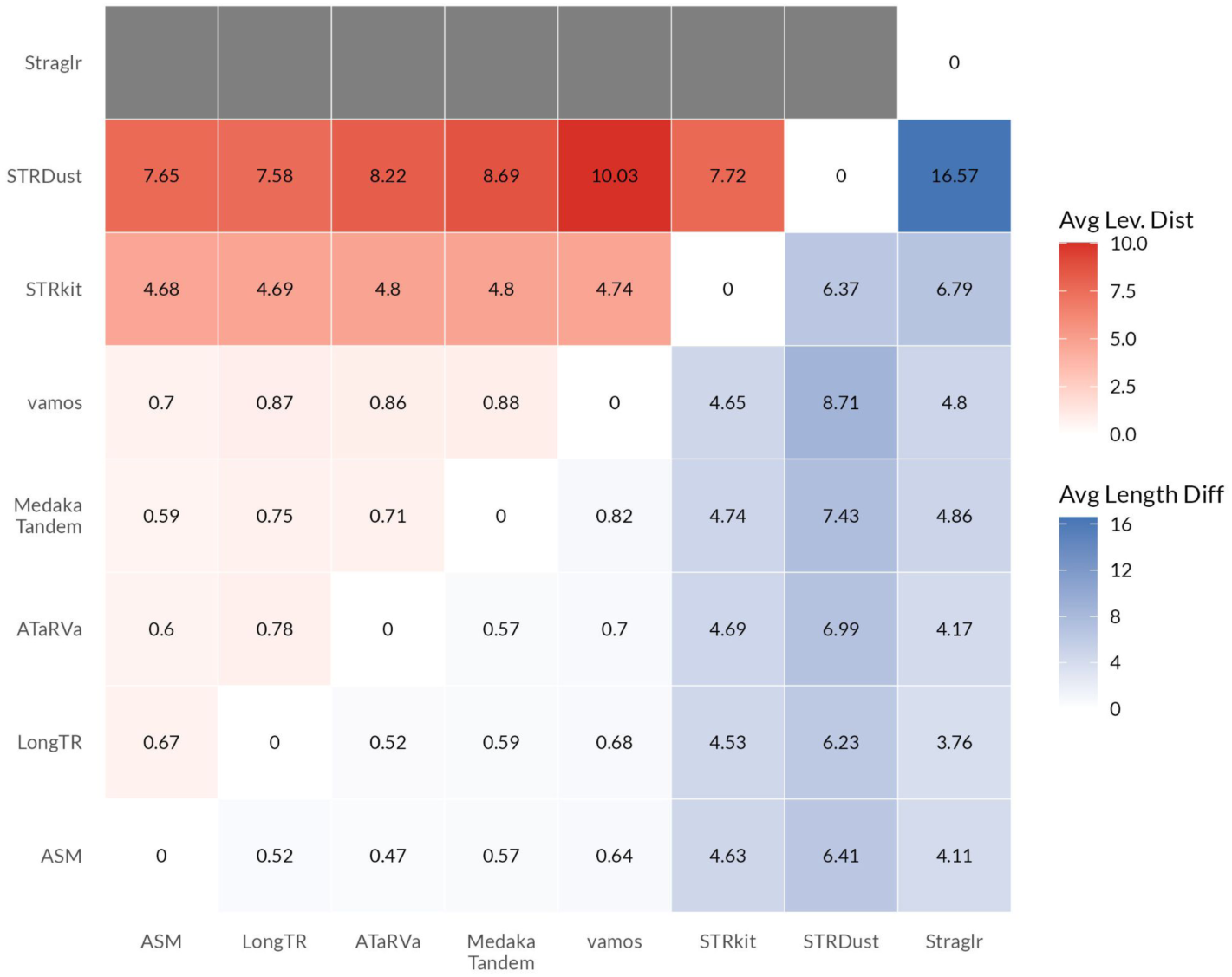
Genotyper consistency: pairwise comparison of genotyping tools based on allele sequence and length differences. The heatmap displays pairwise comparisons between all tools, including genome assemblies as a reference genotyping method. The upper triangle represents the mean Levenshtein distance between allele sequences, and the lower triangle represents the mean absolute length difference between alleles, averaged across samples at only loci genotyped by all genotypers. Straglr is excluded from sequence-level comparisons as it does not report allele sequences.

We next examined loci where the majority of tools disagreed with the assembly calls. Using one representative sample (HG00290), we selected all loci where at least five tools disagreed with the assembly (discordant loci: 402) and visualized pairwise tool comparisons restricted to these loci as a heatmap (**Supplementary figure 9**). At these discordant loci, both sequence and length differences between tool pairs are substantially smaller than the differences observed between individual tools and the assembly. This suggests that either all tools are systematically missing the assembly-based genotype, or that the read data presented to the tools differs from the sequencing data underlying the assembly, leading all tools to converge on the majority genotype supported by the available reads.

#### Sensitivity to Pathogenic Expansions

To assess sensitivity to pathogenic expansions, we ran all tools on blood-derived sequencing of individuals with a known TR disorder and/or an orthogonal molecular test. This previously published study [36] generated targeted ONT reads using adaptive sampling of 37 target genes with known TR expansions. Of those, nine genes had pathogenic expansions in this cohort: *ATXN1*, *C9orf72*, *DMPK*, *FMR1*, *FXN*, *HTT*, *NOTCH2NLC*, *PABPN1*, and *RFC1*. We analysed 27 individuals who had known TR disorder and/or allele-size estimates from molecular testing using RP-PCR, Southern blot, or both **(Supplementary Table 8)**. The cohort included 19 individuals with a molecularly confirmed pathogenic TR expansion, 2 individuals with premutations, 2 carriers, and 3 unaffected family members without expansions. An additional sample was listed as RFC1 positive and clinically affected, but no molecular testing results were available.

Reads from these individuals were aligned to hg38 and genotyped with all 7 tools using the same pipeline as above. We then subsetted the results to only the relevant disease locus for each sample. LongTR did not genotype any known TR disease loci in any of the samples because it requires at least 10 spanning reads, each with a mean base quality of 30. As this is R9 ONT data published in 2022, the mean base qualities per sample were 19.5 (range 18.5-20.8), and no disease loci passed these filters. Other tools genotyped most loci; however, because tools differed in their default read-depth requirements, to maximize sensitivity, we reran all tools and, where possible, adjusted parameters to call any locus with at least one read. The difference was most notable for LongTR (0 to 87%, **Figure 6A**), but most tools increased in sensitivity with these setting adjustments. The exception was Straglr, which detected an additional pathogenic expansion in default settings than in “min reads” settings. The initial analysis was performed with vamos v2.1.7; however, v3.0.5 was released during the analysis phase of this study. We noted a dramatic drop in sensitivity between versions from 61% to 30%, with the missed loci not included in the VCF. v2.1.7 was used for the remainder of the pathogenic loci analysis, and the potential bug has been reported. Vamos changed the read consensus algorithm between versions from abPOA to sPOA. Vamos v3.0.5 results are retained here for transparency and consistency with the assembly and Mendelian analyses, where it performed well.

**Figure 6:**
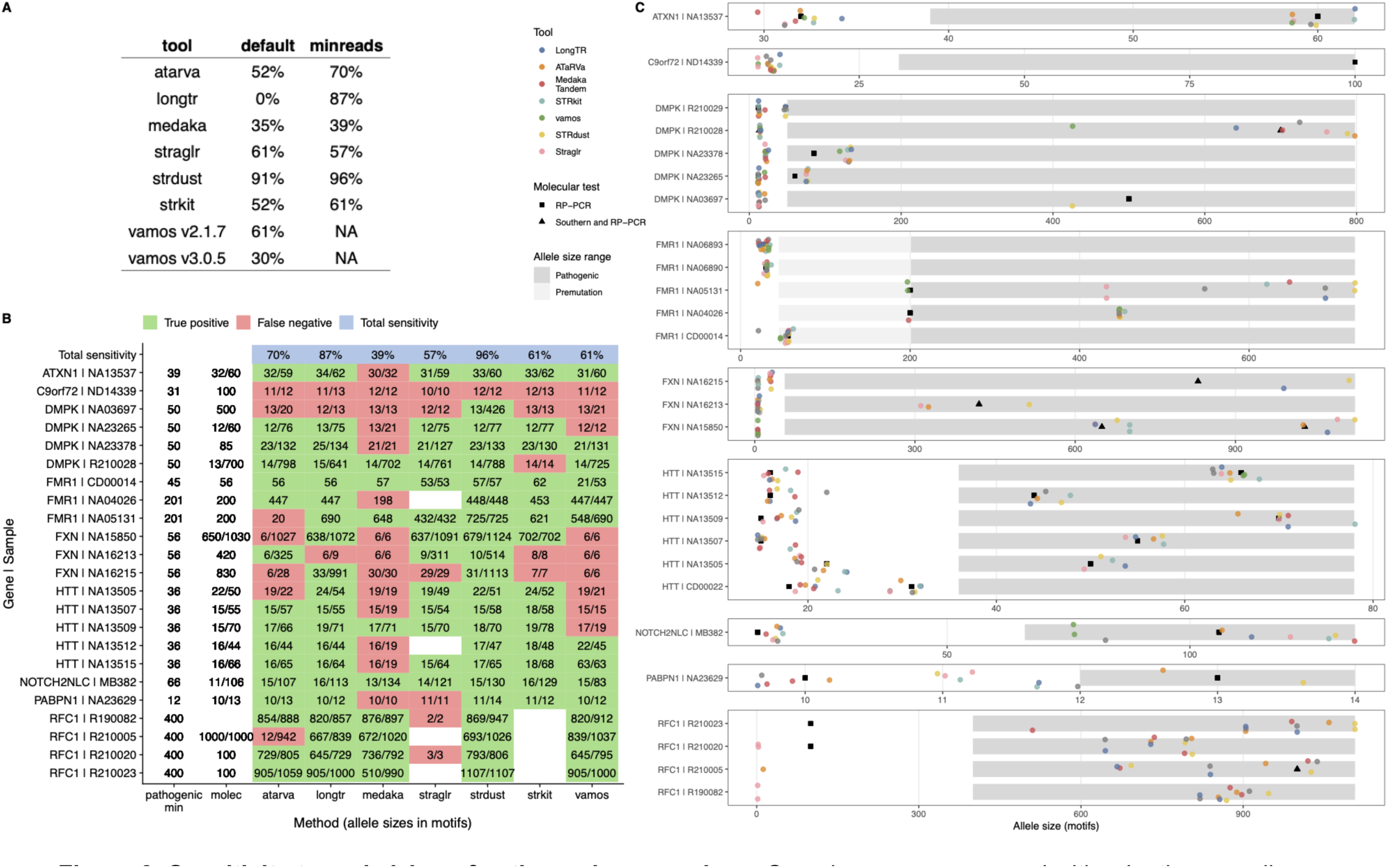
Sensitivity to and sizing of pathogenic expansions. Samples were sequenced with adaptive sampling [36]. Where possible, tools were run with the most sensitive settings to include lower-quality reads, except for Straglr, which was run in default settings. vamos v2.1.7 was used. **A**: Most tools were more sensitive to pathogenic expansions when settings were adjusted to include lower-quality reads. Relevant settings were identified in all tools other than vamos, specifically: ATaRVa (--min-reads 1), LongTR (--min-reads 1 --min-mean-qual 1), Straglr (--min_support 1 --min_cluster_size 1), STRkit (--min-reads 1 --min-read-align-score 0.1), STRdust (--support 1), Medaka Tandem (--min_depth 1). LongTR made no calls in these samples when run with default settings, as no reads passed filters. **B.** Sensitivity to detect pathogenic or premutation alleles based on length. In addition to the individual tool genotypes, the pathogenic min threshold from STRchive is reported, and the size estimates from RP-PCR or Southern blot are indicated (“molec”). Where possible, tool filters were adjusted to consider reads of all qualities and alignment scores. Unaffected individuals were excluded. **C.** Length concordance between RP-PCR and Southern blot molecular assays and genotypes for samples with known pathogenic TR expansions and their unaffected family members.

None of the tools achieved 100% sensitivity (**Figure 6B**), and all missed the *C9orf72* expansion. There was read evidence of the expansion, but the aligner softclipped these reads, so none fully spanned the expansion (**Supplementary Figure 10**). STRdust demonstrated consistently superior sensitivity across settings (91% in the default setting and 96% in the “min reads” setting). While the default settings of LongTR meant it did not call variants at any of these loci due to the lower quality of R9 data, it performed almost as well as STRdust after parameter adjustment (87%).

For long pathogenic alleles such as *FMR1* and *RFC1*, RP-PCR is known to dramatically underestimate allele sizes. We observed much longer sequence-based size estimates for these loci (**Figure 6C**), demonstrating the value of sequencing for sizing large expansions. The genotypers also identified the expected primary motif for all pathogenic loci, although the allele sequences contained numerous interruptions, likely due to high per-base error rates in the sequencing data.

#### Final verdict: no best tool

No single genotyper was universally optimal in our analysis. We benchmarked all tools on performance and accuracy, summarizing the results in a comparative table and a heatmap (**Figure 7**) to help users select a tool suited to their project goals.

**Figure 7:**
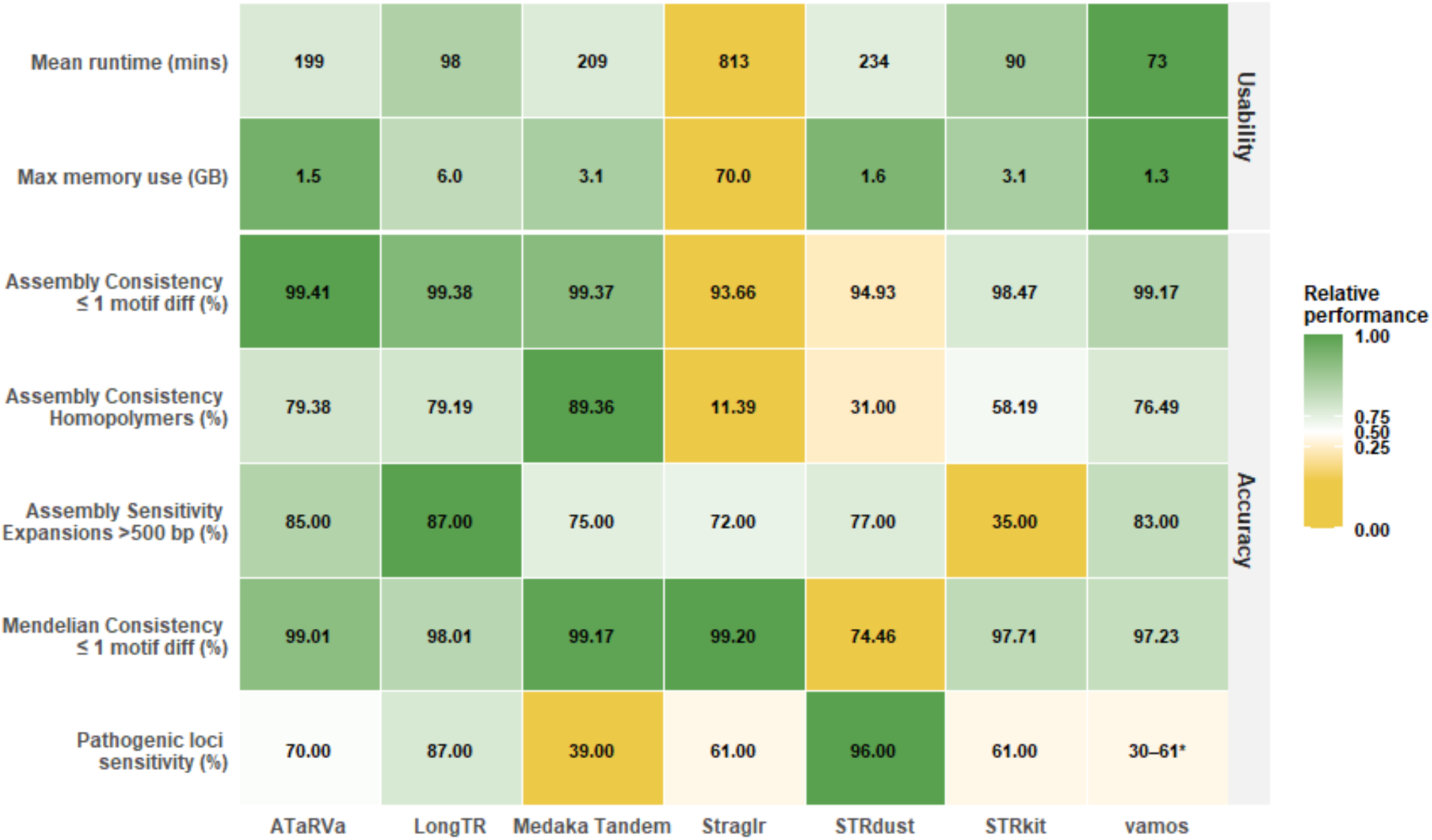
Relative performance heatmap across key metrics of tandem repeat genotyping tools. Cell values show the observed metric, and colors represent performance scaled within each metric (0–1), with darker green indicating better relative performance and yellow indicating worse relative performance. Usability metrics were based on the 43k catalog in GIAB samples, single-threaded (see Methods). Assembly accuracy metrics comprise ≤1 motif unit consistency, homopolymer consistency, and sensitivity for expansions >500 bp. Mendelian consistency represents the proportion of loci consistent with parental genotypes (MATCH + ONE-OFF; ≤1 motif unit difference) across all child genotypes in the HG002 trio. Pathogenic sensitivity: percentage of pathogenic or premutation genotypes identified with the best settings or version. * 61% for vamos v2.1.7, 30% for vamos v3.0.5.

Tools were evaluated on computational performance based on runtime and memory requirements. With the exception of Straglr, all tools ran within configurations typical of personal computers. Vamos stood out with the most favorable resource usage across both metrics.

Tool accuracy was evaluated across three measures: concordance with high-depth assemblies, Mendelian concordance, and sensitivity to pathogenic expansions. No single tool performed best across all assessments, reflecting both differences in tool design and the inherent challenges of tandem repeat genotyping, where error rates vary considerably by locus characteristics (**Figure 7**).

In assembly concordance ATaRVa, Medaka Tandem, and vamos exceeded 99% accuracy. Homopolymer repeats were a consistent challenge across all tools and a major source of inaccuracy, accounting for nearly half of all incorrect calls in the top-performing tools. Medaka Tandem performed substantially better than other tools at homopolymer loci. Large expansions relative to the reference were also challenging for many genotypers. For loci with expansions exceeding 500 bp, LongTR performed best, followed closely by ATaRVa and vamos. For additional discussion of the overall performance of the tools in the context of their algorithmic differences, see **Supplementary Note**.

For Mendelian consistency, Straglr, Medaka Tandem, and ATaRVa all exceeded 99%. A Mendelian consistency of 99% would imply approximately 300-400 apparent *de novo* variants. This likely reflects genotyping error rather than true *de novo* variation.

STRdust emerged as the most sensitive tool for pathogenic expansions, despite its comparatively lower accuracy in genome-wide benchmarking. This discrepancy may be attributable to STRdust’s development, specifically for the analysis of pathogenic variants at clinically relevant tandem repeat loci in ONT [30], and is therefore optimized for this specific use case. Among the remaining tools, LongTR showed the next best performance, which may be attributed to its strategy of retaining all reads (See **Supplementary Note**).

Straglr consistently underperformed across most measures, except Mendelian consistency. This, coupled with it not reporting allele sequencing in the VCF, suggests it should be reconsidered as the default choice in popular workflows such as the EPI2ME wf-human-variation pipeline [42].

LongTR strikes a good balance between accuracy on typical-sized alleles and sensitivity to pathogenic expansions, making it a promising general-purpose choice. However, this result was achieved on R9 data only by reducing read quality and alignment thresholds. In all cases, our initial experimentation suggests that many tools could perform substantially better with further parameter tuning.

## Discussion

In this study, we conducted the first comprehensive sequence-level benchmark of long-read TR genotypers in Oxford Nanopore data. We assess seven genotypers across ∼43k TR loci in more than 100 human genomes. While previous evaluations were typically performed in support of a specific new tool and focused primarily on allele length, we show that a sequence-level approach more accurately estimates accuracy and reveals sequence-level discrepancies, especially for long alleles, VNTRs, and non-reference alleles. By integrating four complementary accuracy metrics, assembly concordance, Mendelian consistency, cross-tool agreement, and sensitivity to confirmed pathogenic expansions, we provide the most complete assessment to date of the strengths and limitations of current long-read TR genotyping tools. We also provide feedback on the user experience, including tool installation, computational resource requirements, and documentation, giving potential users a more realistic idea of what they are getting themselves into when they select a genotyper. Our work closes a critical gap and provides guidance for selecting TR genotypers for population studies, mechanistic research, and clinical diagnostics.

### Trade-offs Between Tools: Is There a “Best” Genotyper?

No single genotyper was universally optimal in our analysis. We identified several excellent options; however, the specific choice depends on the study goals. For example, we observed variability in performance across homopolymers, long alleles, heterozygous loci, and non-reference alleles, reflecting distinct algorithmic sensitivities and error modeling strategies. Accuracy also depended on sequencing chemistry; while all algorithms performed better on higher-quality R10 data, the difference in performance between R9 and R10 was greater for some tools. For example, Medaka Tandem likely performs better on R9 and at homopolymers because it adapts its error modelling on a per-sample basis, effectively handling high-error reads.

Computational performance varied widely, with dramatic differences in memory and run-time, but importantly, we did not observe a consistent trade-off between computational cost and accuracy. Many of the fastest tools were also among the most accurate. Consequently, the “best” tool is context-specific: clinical diagnostic prioritizes sensitivity to very large pathogenic expansions, while population-scale studies require computationally efficient, scalable methods. While all studies are likely to benefit from complete and accurate allele sequences, studies focused on motif evolution will especially benefit from motif decomposition, which is supported by only a small number of tools (e.g., vamos and ATaRVa).

### Strengths and Limitations of Each Benchmarking Approach

Each of our benchmarking strategies offers district and often complementary strengths and limitations, so interpreting tool performance requires balancing what each can reveal. Assembly concordance provides a high-quality, but not infallible, comparison point: we do not assume assemblies are the perfect truth, but rather treat them as deep-coverage, multi-technology, orthogonal gold standards that complement our ∼30x read-based genotyping. Because most TR loci are invariant, tools biased towards calling the reference allele can still score high in assembly length concordance. Mendelian consistency has similar pitfalls: a genotyper that always outputs the reference allele at every locus would achieve perfect Mendelian consistency and still appear to perform relatively well against assemblies, while completely failing to detect pathogenic expansions. However, a tool with high Mendelian consistency may generate fewer false-positive *de novo* variants in a trio analysis. Cross-tool agreement likewise does not necessarily indicate correctness; however, patterns of agreement and disagreement can be informative. When multiple tools agree with each other but diverge from the assembly, this often indicates assembly errors or allelic dropout in the reads, whereas widespread disagreement typically highlights difficult or unstable loci. Finally, sensitivity to pathogenic alleles is the most clinically relevant measure, and our analyses show that tools may perform well under other benchmarking characteristics but still miss many pathogenic expansions, some of the most challenging alleles to genotype. Taken together, these complementary strategies expose different and often non-overlapping error modes, underscoring the importance of multidimensional evaluation rather than reliance on any single metric.

### Usability Challenges: A Major Barrier to Adoption

A major theme emerging from this work is the substantial difficulties we experienced using current long-read TR genotypers in practice. Despite our team of experienced bioinformaticians and the developers of several long-read TR genotyping methods, we spent an outsized portion of this project installing and debugging tools, interpreting cryptic error messages, repeatedly rereading incomplete or outdated documentation, and, in some cases, reading source code or contacting tool authors when documentation gaps became insurmountable. Across tools, we encountered installation failures due to dependency issues, unexpected transformations of genomic coordinates between the input catalog and output VCF (when the tool deemed the catalog coordinates to be too narrow), VCF outputs that required downstream correction or reconstruction, and a general lack of examples, tutorials, or guidance on parameter optimization. These challenges were not limited to newer or more niche tools; even widely used methods presented similar obstacles. Given our own experience, we expect that a general bioinformatics user will face even greater barriers to adopting these tools, threatening both usability and reproducibility. We advocate for improved documentation, standardizing output formats beyond the current VCF specification, providing clearer error messages, and offering users practical guidance on parameter selection and workflow integration. These changes would dramatically enhance the accessibility and reliability of TR genotyping methods and accelerate their uptake across genomics from population studies to clinical settings.

### Parameter Optimization: Essential but Under-documented

Parameter optimization emerged as another critical but under-documented aspect of TR genotyping. For the majority of our benchmark analyses, we deliberately focused on default or recommended parameters so as to reflect real-world usage. However, many tools expose dozens of tunable parameters that can substantially affect accuracy, sensitivity, and runtime. In our preliminary explorations of pathogenic loci, we observed dramatic improvements in sensitivity for most tools when parameters were adjusted to include lower-quality and less well-mapped reads. This suggests that our benchmark likely underestimates the maximum achievable performance of several tools. Effective parameter tuning is likely essential for specific scenarios, including long (i.e., most pathogenic) alleles, loci with significant somatic mosaicism, and regions dominated by homopolymers. Such optimization is likely to be especially critical in clinical settings to ensure sensitivity. Yet, guidance on parameter selection is rarely provided in tool documentation, nor explanations of how different settings interact with sequencing chemistry, coverage, or other error profiles. Although comprehensive parameter optimization was beyond the scope of this benchmark, our findings suggest some starting points for pathogenic loci and a call for developers to prioritize more comprehensive guidance and best-practice recommendations.

### Towards an Ideal TR Genotyper

Although several tools performed well across particular scenarios, no single genotyper emerged as universally optimal, underscoring the need for continued method development. An ideal TR genotyper would offer practical usability, easy installation, clear documentation, and a standardized, interpretable VCF format. This should be paired with robust sequence-level performance across STRs and VNTRs. It should accurately reconstruct full allele sequences, resolve complex multi-motif structures, and maintain high accuracy across long alleles, non-reference alleles, and homopolymer-rich regions. Computational efficiency, explicit modeling of sequencing error, and optional read-level information to assess somatic variation would further enhance utility. Features such as haplotype phasing, uncertainty estimates, and built-in motif decompositions would make such a tool broadly applicable to population and clinical studies. Our findings highlight the methodological and usability gaps that must be addressed to approach this ideal and provide clear priorities for our community to enhance TR genotypers.

### Strengths and Limitations of This Study

This study has several strengths. It is the first large-scale, sequence-level benchmark of long-read TR genotypers, conducted independently of any single tool. It included multiple orthogonal accuracy metrics, pathogenic expansions, a catalog enriched for challenging loci, and an independent assessment of usability and computational performance supported by reproducible workflows. Nonetheless, several limitations should be noted. Several authors have contributed to tools benchmarked in this study, specifically ATaRVa, STRdust, and vamos. We attempted to mitigate some of this bias by contacting the authors of all the tools we benchmarked via their respective GitHub repositories when we encountered issues. Another limitation of our approach is that we predominantly used default parameters, and our limited exploration of this space indicates that default settings may underestimate performance, especially for higher-error data and pathogenic alleles. We did not assess the impact of somatic mosaicism on tool performance, nor the ability of tools to detect it. We also did not benchmark special features specific to a small number of tools, such as motif decomposition. Finally, this benchmark represents a snapshot of a rapidly evolving field, and we hope many of the tool limitations we observed will be addressed in future versions.

## Conclusions

Our results show that no single genotyper performs best across all loci, metrics, and conditions; however, several tools offer strong performance for specific applications, while others provide more consistent performance across metrics, providing researchers with a set of good options rather than a single perfect solution.

This benchmark advances the field in several important ways. It demonstrates that sequence-level accuracy allows for more differentiation between genotypers than length accuracy alone. It highlights conditions under which accuracy deteriorates across many tools: homopolymers, higher sequencing error rates, and large pathogenic expansions. The benchmark highlights key methodological and usability challenges and calls for further development of both the tools themselves and the documentation for optimizing them.

Beyond evaluating current methods, this work offers practical recommendations to guide tool selection based on study goals and provides fully reproducible workflows to support community reuse, transparency, and future method development. Together, these contributions lay a foundation for the next generation of large-scale TR analyses, including those undertaken by the 1000 Genomes Long-Read Sequencing Consortium, and strengthen the broader scientific groundwork for studies of TR mutation dynamics, population variation, and clinical diagnostics.

## Methods

### Selection and prioritization of tools for TR benchmarking

In the initial phase, we reviewed the literature to identify software tools developed for TR genotyping. The primary source of TR methods was the long-read-tools.org resource [38,39], which was supplemented by searching Google Scholar for references to TR genotypers, including those in methods sections of analysis papers. We curated key characteristics, including supported sequencing technologies, handling of compound repeat structures and non-canonical motifs, detection of novel loci, and the nature of the output (haplotype-resolved genotypes, full allele sequences, and supported motif sizes), which yielded a list of 25 tools (**Supplementary Table 1**). For benchmarking, we selected TR genotypers that report full allele sequences and are under active development (at least one update within the year preceding 11 November 2025). We also added the most popular length-only genotyper, straglr, for comparison. The final selection included 8 tools. (**Supplementary Table 2**).

### Selection of the catalog for benchmarking

The TRExplorer catalog is a comprehensive TR catalogue built from merging individual catalogs, including perfect TRs, adotto benchmark loci, polymorphic loci, VNTR catalogs, and disease-associated TR loci [37]. We used Version 2 of the TRExplorer catalogue, which comprises 5.5 million loci in the human genome, of which ∼100k are variation clusters: regions with closely spaced polymorphic TR loci. This catalog is large in terms of the time required for genotyping individual tools and also poses tool-specific problems during reformatting. For the first phase of benchmarking, we limited our analyses to a subset of loci that were non-overlapping to reduce genotyping complexity. Specifically, we selected non-overlapping loci separated by a minimum distance of 25bp to obtain a set of isolated TR loci. We then sampled 1000 loci for each motif size in the set. We ensured that disease-associated loci were included by adding the STRchive loci [5] back into our testing catalog. This gives us a total of 43,009 loci.

### Sequencing data preparation

The ONT sequencing data for the seven GIAB samples (HG001-7) were downloaded from EPI2ME’s Genome in a Bottle Data Release 2025.01. Each sample is sequenced on 2 flow cells, and the data is available as aligned Compressed Reference-oriented Alignment Map (CRAM) files. One CRAM file for one flow cell run of the sample, hence 2 CRAM files for a sample. For each sample, the data was pooled across both flow cells and downsampled to near 30x coverage. The CRAM files of both the flow cells for each sample are merged using samtools merge followed by samtools view -s 42.{fraction} for downsampling to 30x.

For the HPRC cohort, using the HPRC Data Explorer [48], we selected ONT runs expected to have ∼30-35x sequencing depth in a single flow cell. We balanced samples sequenced on R9 and R10 chemistry: 33 R9.4.1 chemistry and 34 R10.4.1 for a total of 67 samples:

HG00290 HG01786 HG02630 HG04228 NA20129 HG00320 HG01975 HG02698 NA18505 NA20282 HG00321 HG01993 HG02723 NA18508 NA20346 HG00329 HG02004 HG02818 NA18608 NA20762 HG00350 HG02040 HG02922 NA18620 NA20806 HG00408 HG02071 HG02984 NA18948 NA20809 HG00597 HG02132 HG03017 NA18952 NA20870 HG00609 HG02135 HG03209 NA18960 NA21093 HG00639 HG02280 HG03453 NA18974 NA21106 HG01081 HG02293 HG03521 NA19240 NA21110 HG01255 HG02392 HG03710 NA19682 NA21309 HG01346 HG02572 HG03804 NA19776 HG01433 HG02583 HG03816 NA19835 HG01496 HG02602 HG04157 NA19909

ONT reads were extracted using samtools and aligned to hg38 using minimap2 (see https://github.com/dashnowlab/TR-Benchmarking/blob/main/align_HPRC/main_align.nf for more details).

~~~
samtools fastq -T MM,ML ${unaligned_bam} | \
minimap2 -a -y -t ${threads} -R “${readgroup}” ${ref} - | \
samtools sort -@ ${threads} -O CRAM --reference ${ref} -T
${sample}.tmp -o ${sample}.cram
~~~

Coverage estimation was done using mosdepth v0.3.10 [49] on each alignment file.

### Implementation and execution of selected TR genotyping tools

All benchmarking workflows were implemented using the Nextflow workflow management system (DSL2), enabling scalable and reproducible execution across high-performance computing environments. Each Nextflow process was defined with a fully self-contained environment, avoiding reliance on inherited shell modules and preventing dependency conflicts to ensure consistent behavior across compute nodes. Tool dependencies were managed using either Singularity container images or Conda environments, depending on software compatibility: mosdepth, LongTR, Straglr, and STRkit were deployed via containers (Wave/Seqera community images or GitHub Container Registry), whereas ATaRVa, Medaka Tandem, and vamos required dedicated Conda environments; STRdust was executed as a statically pre-compiled binary. Several software bugs and compatibility issues were encountered during implementation and were reported to the respective tool developers. Eight TR genotypers (ATaRVa, LongTR, Medaka Tandem, Straglr, STRdust, STRkit, TRsv, and vamos) were evaluated using a catalogue of approximately 43,000 loci. We successfully generated results for seven tools. TRsv was excluded from downstream analyses due to persistent and reproducible runtime errors originating from its distributed Perl scripts and the Singularity container provided in the authors’ repository, which prevented successful execution on our datasets. The final benchmarking pipeline was applied to 7 GIAB samples, 67 HPRC samples, and 31 samples from the Stevanovski 2022 cohort [36].

#### ATaRVa

ATaRVa (v0.5.0) was run using a minimum of one supporting read per allele (--min-reads 1) parameter and passed the sample karyotype.

#### LongTR

We evaluated LongTR (v1.2) using --alignment-params -1.0, -0.458675, -1.0, -0.458675, - 0.00005800168, -1.0, -1.0, thereby increasing the match-to-insertion and match-to-deletion transition probabilities relative to the default settings to better accommodate the indel-rich error profile of long-read sequencing at TR loci as recommended by the authors. Prior to calling, we computed the maximum TR length in the locus catalogue and passed it as the --max-tr-len parameter to ensure that LongTR could accommodate the longest repeat tract in our benchmark catalog. For samples with XY karyotype, chrX and chrY were treated as haploid by passing `--haploid-chrs chrX,chrY`.

#### Medaka Tandem

We used the Medaka (v2.1.1) tandem-repeat calling module: Medaka Tandem (tandem.py, https://github.com/nanoporetech/medaka/tree/master/medaka/tandem). Sample sex was passed to Medaka as `male` or `female` based on the sample karyotype. For CRAM files that had been aligned in a different environment, Medaka runs failed with htslib cram_get_ref errors because the CRAM header UR tag pointed to an obsolete reference FASTA path, causing the underlying library to attempt to load a non-existent reference despite the correct GRCh38 FASTA being supplied on the command line. We had to re-header these samples to produce Medaka output.

https://github.com/nanoporetech/medaka/issues/574

#### Straglr

Straglr (v1.5.5) was executed using a Python wrapper, with alleles genotyped by size in base pairs (--genotype_in_size). Sample sex was passed as m or f based on sample karyotype.

#### STRdust

STRdust (v0.24.2) was run requesting unphased genotypes (--unphased). For samples with karyotype XY, we passed --haploid chrX,chrY so that STRdust treated chrX and chrY as haploid, while autosomes in all samples were modelled as diploid.

#### STRkit

STRkit (v0.24.2) was run in targeted calling mode with ploidy passed per sample (--ploidy) and sex chromosome karyotype supplied via --sex-chr. Local realignment around TR loci was enabled (--realign).

#### Vamos

vamos (v3.0.5) and (v2.1.7) were run using read-based calling (--read) with the addition of -S to include allele read consensus sequences in the VCF output. These consensus sequences reflect the raw output of the read consensus algorithm, prior to motif set interpretation. Sample allele sequences are reported in the VCF in a non-standard format, appended to the end of the sample field. The reference allele is always “N”, and all GTs are 1/1 or 1/2; therefore, to use this VCF with downstream tools, we extracted the reference allele sequence from the reference genome using the locus start and end coordinates, compared sample alleles to the reference, and marked perfectly matching alleles as “0”. Any non-reference alleles were moved to the ALT field. This post-processing was implemented in a custom fix-vcf.py script, which must be adjusted for the two versions of vamos separately to produce valid VCF format for downstream analysis.

To evaluate sensitivity to pathogenic repeat expansions, all seven tools were additionally run with minimum supporting-read thresholds. In this configuration, ATaRVa and LongTR required at least one supporting read (--min-reads 1), LongTR additionally applied a minimum mean base quality of 1 (--min-mean-qual 1), Medaka used a minimum depth of 1 (--min_depth 1), Straglr required at least one supporting read (--min_support 1) with a minimum cluster size of 1 (--min_cluster_size 1) and a maximum of two clusters per locus (--max_num_clusters 2), STRdust required at least one supporting read (--support 1), and STRkit required at least one supporting read per locus (--min-reads 1). All other parameters remained identical to the standard runs described above. Vamos was run with identical settings to the standard run, as it does not expose a minimum read support parameter.

### Extracting allele information from VCFs

Each tool uses slightly different INFO and FORMAT fields in the VCF. For tools that encode alleles as explicit nucleotide sequences in the REF and ALT fields (all except Straglr), allele sequences for each sample were resolved by mapping the GT indices to the corresponding REF or ALT entry, and loci with missing or unresolvable genotypes were excluded. For Straglr, which does not report explicit allele sequences but instead uses symbolic ALT entries, allele lengths were derived directly from INFO field annotations. The reference allele length was obtained from the RUL_REF INFO field, where available, with sequential fallback to END – POS + 1. ALT allele lengths were extracted from the RB INFO field, which records the repeat block length in base pairs for each alternative allele, and GT indices were used to assign the two allele lengths per sample.

### Benchmarking against assembly

Phased HPRC assemblies were aligned to the GRCh38 reference genome using minimap2 with the -x asm5 preset to generate BAM files. Allele lengths at each locus were calculated by lifting over locus coordinates from the reference to the sample assemblies using these alignments, implemented in an in-house Python script (assembly_concordance.py; https://github.com/dashnowlab/TR-Benchmarking/scripts/assembly_concordance.py).

Some tools adjust the coordinates for a subset of input TR loci. STRkit scans the flanking sequences of each TR locus in the reference for possible repeat extensions and extends the coordinates if matching motifs are found. LongTR expands locus coordinates to include flanking sequences in which sequence variations are identified. For assembly-based concordance, genotypes are therefore evaluated using tool-specific coordinates by performing liftover on the coordinates as reported in each tool’s output VCF. Only loci with single-contig alignments between the reference and the haploid assemblies were retained, and diploid genotypes were constructed by calculating allele lengths independently from each haplotype.

Allele sequences or allele lengths (in the case of Straglr, which does not report sequences) were extracted from each tool’s VCF (See Extracting allele information from VCFs), and the corresponding assembly alleles were extracted based on the VCF coordinates. VCF alleles were then paired with assembly alleles to minimize total pairwise deviation. For tools reporting allele sequences, deviation was calculated as Levenshtein distance; a measure of the minimum number of single-nucleotide edits (insertions, deletions, or substitutions) required to transform one sequence into another. For Straglr, deviation was calculated as the absolute difference in allele length. Length deviations were categorised into four concordance classes: Match (no deviation), Off by 1 bp (deviation of exactly 1 bp), Off by motif (deviation ≤ the motif length of the locus), and Mismatch (deviation exceeding the motif length). For each sample and tool, this procedure generates a per-locus concordance report containing the tool-called and assembly-called alleles, the length deviation, the Levenshtein distance, and the concordance class. This report serves as the basis for all subsequent assembly concordance analyses.

### Mendelian consistency

To assess the accuracy of tandem repeat genotyping, we evaluated Mendelian consistency across the parent–offspring HG002 trio. HG002 (the offspring) is male (46, XY); because loci on the X and Y chromosomes are hemizygous in the offspring, sex chromosomes were excluded, and Mendelian consistency was assessed exclusively on autosomes. Alleles for each sample in the trio are extracted (See Extracting allele information from VCFs). Loci where any trio member had a missing call were excluded. Allele lengths in base pairs were then derived from the resolved sequences, and each individual’s genotype was encoded as a four-dimensional feature vector [L₁, L₂, ΔL₁, ΔL₂], where L₁ and L₂ are the lengths of the two alleles and ΔL₁ and ΔL₂ are their deviations from the reference allele length (Lref = len(REF)). For Straglr, the same four-dimensional feature vector was constructed based on the two allele lengths.

In both scripts, VCFs for the trio members were stream-merged by chromosomal position (CHROM, POS), with LongTR merged by variant ID due to its coordinate representation. Loci present in any subset of trio members were processed only when both parental records were available. Mendelian consistency was assessed by enumerating all four possible parental transmission combinations (one allele from each parent) and both orderings of the offspring’s allele pair, yielding a total of eight candidate pairings. For each pairing, the Manhattan distance between the parental transmission feature vector and the offspring feature vector was computed, and the minimum across all pairings was taken as the locus-level inheritance distance. A locus was classified as Mendelian-consistent (MATCH) if this minimum distance equalled zero, indicating that the offspring’s two allele lengths were exactly explained by one allele from each parent. To capture near-consistent genotypes, we additionally defined three lenient categories: ONE-OFF, where the minimum distance corresponded to a single base pair discrepancy per allele (half-distance = 1 bp); MLEN-OFF, where the discrepancy was within one motif length unit; and MISMATCH for all remaining loci. Analyses were stratified by offspring genotype class (0/0, 0/1, 1/1, and 1/2) and by repeat motif length, with the latter binned into unit categories of 1–10 bp and a “>10 bp” group. Mendelian consistency rates were reported both per genotype class and aggregated across loci, with the combined rate of MATCH, ONE-OFF, and MLEN-OFF referred to as the “good” rate.

### Concordance between genotypers

VCF output files from each tool are compared using the run_comparisons.py script available at https://github.com/dashnowlab/TR-Benchmarking. Some tools change the start and/or end coordinates of loci from those provided as input. To ensure comparisons occur at the same genomic positions across files, the catalog is used as a positional reference. Before comparisons are performed for a given line, each tool locus is assigned to the overlapping catalog position. If a VCF does not contain a matching locus, that tool is skipped until its coordinates align with the catalog again.

Once synchronized, the script first determines if the tools are widening or narrowing the coordinate boundaries relative to the catalog. Finally, the specific alleles from each tool are compared using length differences and the Levenshtein distance (a metric representing the total insertions, deletions, and substitutions needed to transform one allele into the other). Straglr did not provide sufficient information to reconstruct a sequence for Levenshtein distance analysis. Consequently, only length and positional comparisons were performed for that tool.

### Pathogenic loci

We analysed previously published Stevanovski 2022 cohort [36] blood-derived ONT sequencing data from 27 individuals with a diagnosis and/or allele-size estimates from molecular testing using RP-PCR, Southern blot, or both (**Supplementary Table 8**). Targeted ONT sequencing was generated using adaptive sampling of 37 target genes with known TR expansions (see Supplementary Table 1 in [36]), including those with expansions in the samples: *ATXN1*, *C9orf72*, *DMPK*, *FMR1*, *FXN*, *HTT*, *NOTCH2NLC*, *PABPN1*, and *RFC1*.

Raw reads were aligned to hg38 using minimap2 and genotyped with all seven tools using the same pipeline and 43k TR catalog as described above. We ran two versions of vamos, v2.1.7 and v3.0.5, because a new version was released during the analysis phase of this study. We then subsetted the results to only the relevant disease locus for each sample by filtering to VCF rows with a starting position within +/- 50 bp of the disease locus start position in STRchive-disease-loci-v2.15.0.hg38.general.bed (https://github.com/dashnowlab/STRchive/releases/tag/v2.15.0) [5].

The PABPN1 locus has different pathogenic thresholds in the literature: 8 vs 12, depending on whether you include all GCN polyalanine codons or only pure GCG repeats [50]. The RP-PCR validations in [36] used the narrower definition, while STRchive [51] uses the wider definition; therefore, we added 4 motifs to the RP-PCR values to make them consistent with the modern definition.

Tool sensitivity to pathogenic or premutation expansions was assessed by comparing the length of the allele in the tool VCF with the pathogenic threshold (pathogenic min) or, for premutation alleles, the intermediate threshold (intermediate min) from STRchive. If the tool allele length was greater than or equal to the threshold, it was marked as a successful detection. For dominant diseases, one allele above the threshold was required; for recessive diseases, two. For X-linked conditions, if tools reported two alleles, at least one allele above the threshold was required in hemizygous males. Some tools do not accept sample sex as input.

## Data availability

All sequencing datasets analyzed in this study are publicly available and were obtained from reference genomes generated by the Genome in a Bottle (GIAB) consortium. Long-read whole-genome sequencing data for samples HG001–HG007 were obtained from Jan 2025 sequencing data released by EPI2ME of GIAB samples (s3://ont-open-data/giab_2025.01/). HPRC genomes were identified using the Data Explorer (https://data.humanpangenome.org/raw-sequencing-data) and downloaded from their AWS s3 bucket (s3://human-pangenomics/). Adaptive sampling targeted sequencing of samples with pathogenic expansions were downloaded from the NCBI Sequence Read Archive (SRA) under accession no. PRJNA786382. All other data supporting the findings of this study are available either within the article, the supplementary information, or the supplementary data files.

Downsampled and aligned CRAMs and VCFs generated by our study are available at https://s3.amazonaws.com/1000g-ont/index.html?prefix=working_dir/tr_benchmarking/ or https://drive.google.com/drive/folders/1CwxeqE1TWZZ8XklSfn0jVt_fUoOtZJdQ, to be deposited in an appropriate repository prior to publication.

## Code availability

The study utilized previously published analysis tools and custom scripts described in the Methods section. Custom scripts developed for this study are available in a public repository at https://github.com/dashnowlab/TR-Benchmarking and will be archived at Zenodo upon publication.

## Acknowledgements

The authors would like to thank Brent Pedersen for feedback on the early concept and methodology, and Laurel Hiatt for feedback on the manuscript. We are grateful to the other members of the 1000 Genomes Long-Read Sequencing Consortium who provided feedback and support, especially Theo Nelson for feedback on this manuscript and Miten Jain for assistance with data management and sharing.

Finally, we thank the developers of the TR genotyping tools, many of whom were encouragingly responsive, open to feedback, and generous with advice. In particular, Readman Chiu, Helia (Helyaneh) Ziaei Jam, and Sergey Nurk.

## Author information

### Contributions

Contributions are described following the https://credit.niso.org/ framework.

Ebay Aliyev: Conceptualization, Data Curation, Formal analysis, Software, Writing - Original Draft, Writing - Review & Editing, Visualization

Akshay Avvaru: Conceptualization, Data Curation, Formal analysis, Software, Writing - Original Draft, Writing - Review & Editing, Visualization

Harriet Dashnow: Conceptualization, Data Curation, Formal analysis, Software, Writing - Original Draft, Writing - Review & Editing, Visualization, Supervision.

Wouter De Coster: Conceptualization, Writing - Review & Editing

Elizabeth Ostrowski: Conceptualization, Writing - Review & Editing

Denis M Nyaga: Writing - Review & Editing

Danny E. Miller: Writing - Review & Editing

