## Supplementary Note and Figures for "A comprehensive assessment of tandem repeat genotyping methods for Nanopore long-read genomes"

#### Algorithmic basis of tool performance

The overall accuracy differences between tools can be traced to their distinct strategies across four key steps: read selection, repeat identification, haplogroup assignment, and consensus generation.

In read selection, tools apply different filtering criteria that directly affect sensitivity. STRkit applies a stringent mapping quality threshold, filtering out many informative reads and resulting in a high rate of uncalled loci. LongTR applies a high mean Phred quality threshold, which prevents it from making calls on R9.4.1 data where mean basecall quality falls below this threshold. For repeat identification within reads, most tools rely on the original read alignment to define the repeat boundaries. LongTR and STRdust are exceptions, performing local realignment before determining the repeat sequence within each read. STRkit takes a motif-centric approach, aligning reads to perfect iterations of the input motif sequence.

Haplogroup assignment is arguably where tools diverge most substantially. LongTR iteratively builds candidate haplotypes from the reads and assigns reads by aligning them to these candidates. ATaRVa selects the most informative, high-quality SNPs from the reads or applies k-means clustering based on repeat lengths. Vamos segregates reads by max-cut heuristic that initializes partitions with the two reads with the most disagreeing heterozygous SNVs and assigns the remaining reads to a partition based on shared SNVs. STRkit supports multiple phasing strategies, including input haplotags, an external SNV VCF, de novo identification of informative SNVs combined with repeat length, or a Gaussian mixture model approach based solely on repeat lengths. Straglr applies a Gaussian mixture model to segregate reads into haplotype groups based on repeat length distributions. STRdust uses haplotag information from alignment files by default, or performs hierarchical clustering based on pairwise read alignments.

Finally, consensus generation relies on partial order alignment (POA) across all tools that report allele sequences (except Straglr). Specifically, ATaRVa, Medaka Tandem, and vamos use the Python distribution of absolute banded partial order alignment implementation (abPOA, https://github.com/yangao07/abPOA/tree/main [1]), while LongTR, STRkit, and STRdust use C++ and Rust distributions of SIMD accelerated partial order alignment (sPOA, https://github.com/rvaser/spoa [2]). The tools also slightly differ in the choice of reads within a haplogroup for consensus generation. ATaRVa restricts consensus building to reads of allele length equal to the mode repeat length within the haplogroup. STRdust identifies and excludes outlier reads before consensus generation. LongTR additionally applies a scoring model that aligns reads across different haplotype combinations to determine the final genotype. While all tools follow the same broad workflow of read filtering, repeat identification, haplogroup assignment, and consensus generation, it is the specific combination of strategies at each step that drives the observed differences in accuracy across tools and locus types.

Pathogenic locus genotyping presents unique challenges in tandem repeat analysis, as these loci exhibit error profiles and instability not typically observed at other genomic loci. To maximize tool sensitivity at these loci, we applied more lenient thresholds to ensure all available read information was considered. However, accurately calling expansions at pathogenic loci requires appropriately weighting longer alleles, as sequencing coverage of expanded alleles decreases with increasing length, increasing the risk of missed calls. Additionally, the inherent instability of pathogenic loci introduces higher allelic variance, requiring tools to weigh reads appropriately to accurately resolve the expanded allele.

#### Supplementary Note References


1. Gao Y, Liu Y, Ma Y, Liu B, Wang Y, Xing Y. abPOA: an SIMD-based C library for fast partial order alignment using adaptive band. Bioinformatics. 2021;37:2209–11.

2. Vaser R, Sović I, Nagarajan N, Šikić M. Fast and accurate de novo genome assembly from long uncorrected reads. Genome Res. 2017;27:737–46.

### Supplementary Figures

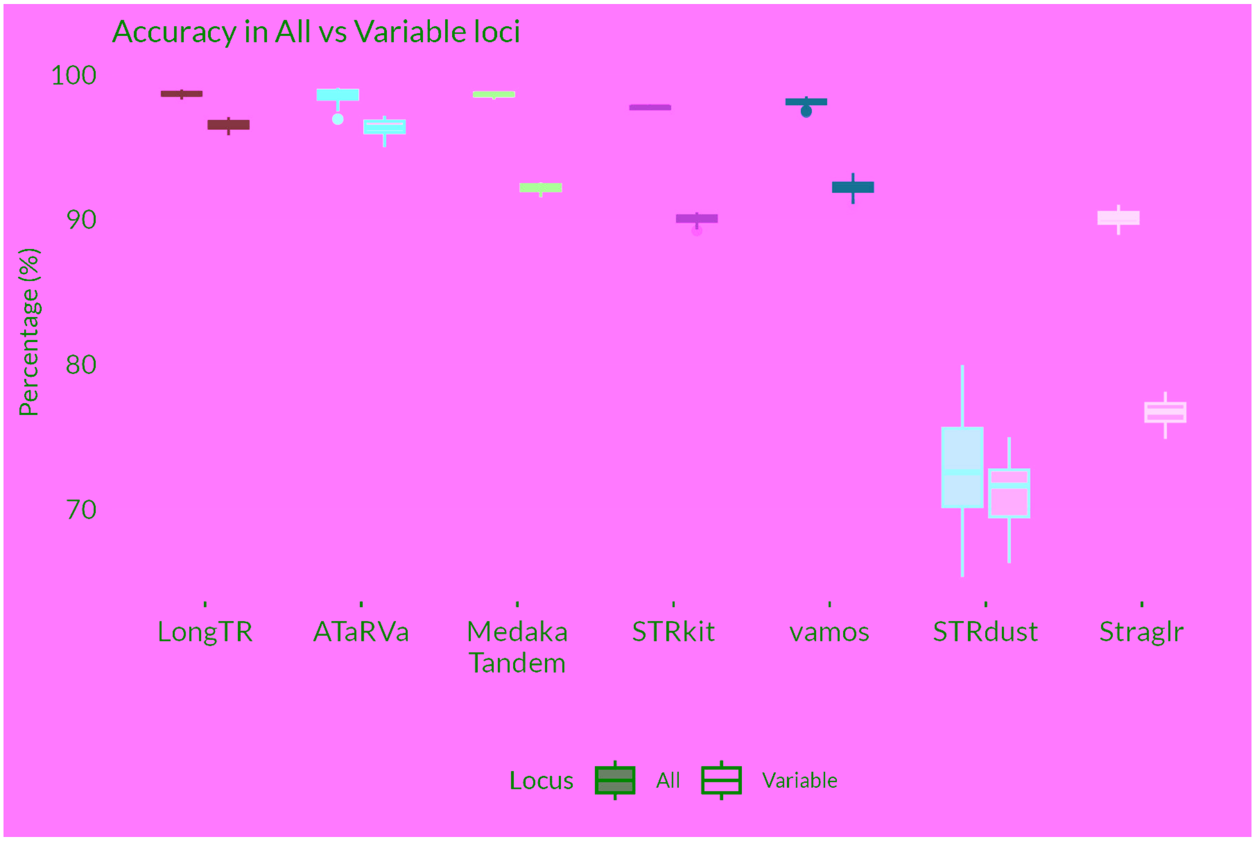


**Supplementary Fig 1**: **Accuracy of tools across all loci and variable loci in R10.4.1 samples.** Box plots display the distribution of accuracy for each tool across samples sequenced with R10.4.1 pore chemistry. Fully opaque boxes represent accuracy calculated across all loci, while transparent boxes represent accuracy restricted to variable loci (excluding homozygous reference calls), allowing direct comparison of tool performance in the presence and absence of non-polymorphic loci.

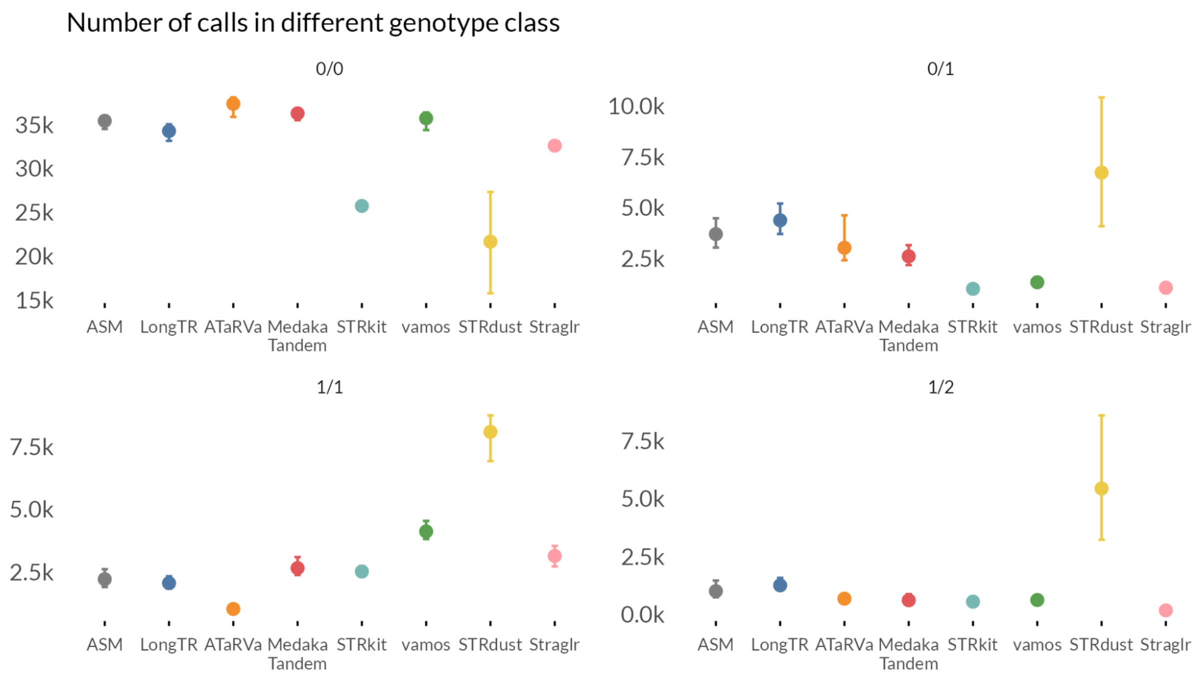


**Supplementary Fig 2**: **Number of loci called by each tool across genotype classes.** Points represent the mean number of loci called by each tool within each genotype class (0/0: homozygous reference, 0/1: heterozygous reference, 1/1: homozygous alternate, and 1/2: heterozygous alternate). Error bars indicate the range between the minimum and maximum number of calls observed across samples within each genotype class.

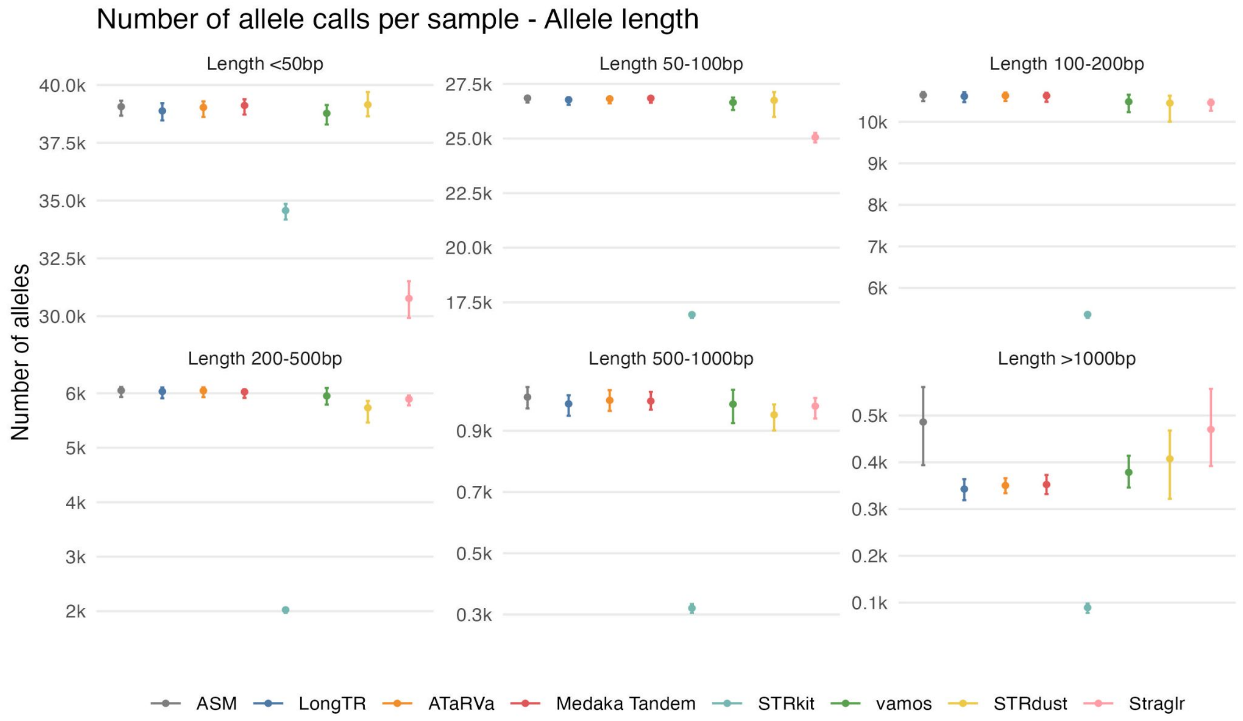


**Supplementary Fig 3**: **Mean number of alleles called per tool across allele length categories.** Points represent the mean number of alleles called by each tool across samples within allele length categories (<50 bp, 50–100 bp, 100–200 bp, 200–500 bp, 500–1000 bp, and >1000 bp). Error bars indicate the range between the minimum and maximum number of calls observed across samples within each category.

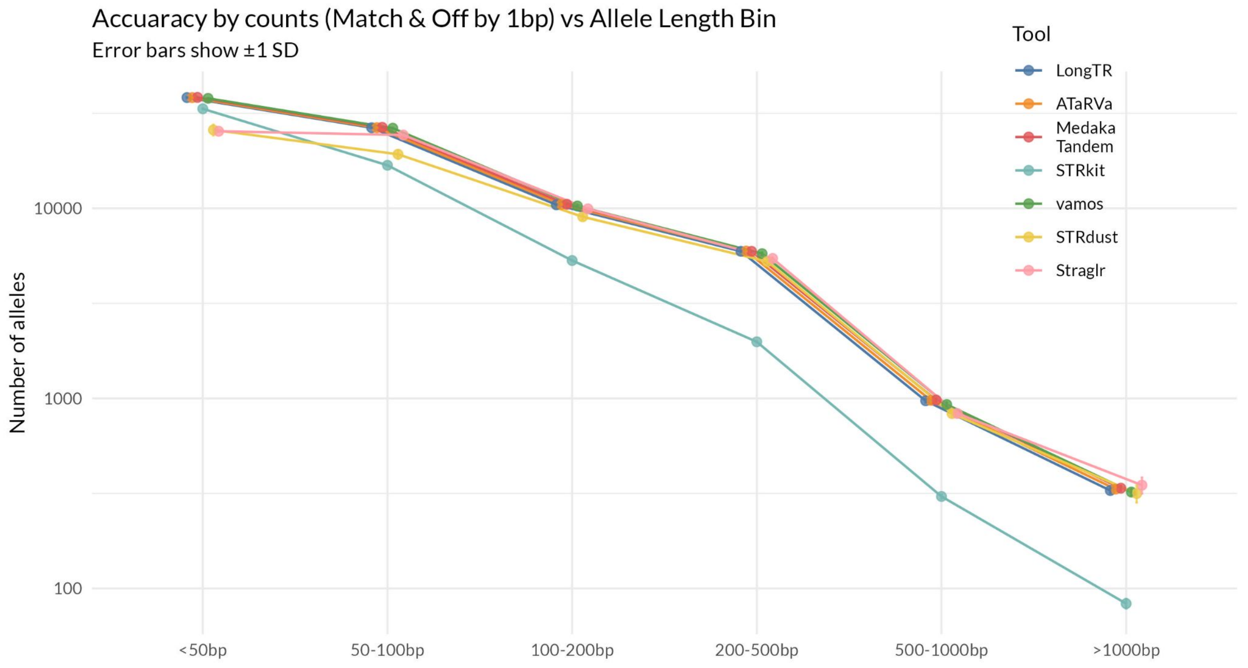


**Supplementary Fig 4**: **Mean number of accurate calls per allele length category across all samples.** Points represent the mean number of accurate calls made by each tool across all samples within allele length categories (<50 bp, 50–100 bp, 100–200 bp, 200–500 bp, 500–1000 bp, and >1000 bp). Y-axis indicates the number of alleles (log scale). Error bars indicate the range between the minimum and maximum number of accurate calls observed across samples within each category.

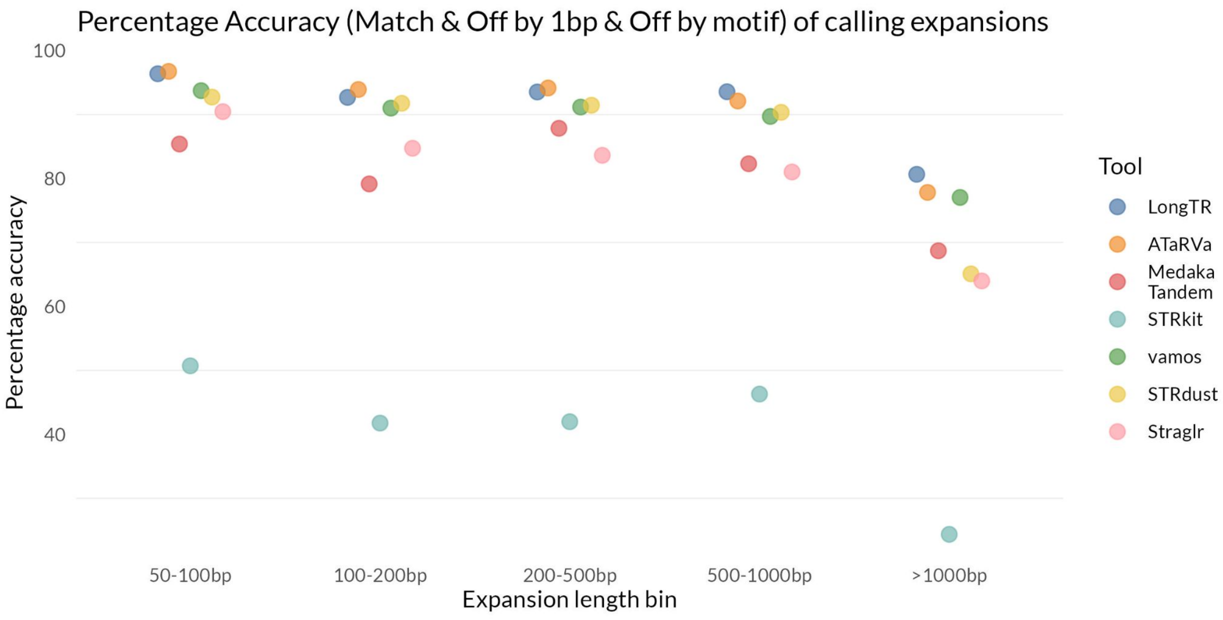


**Supplementary Fig 5**: **Accuracy of expansion calls per tool across R10.4.1 samples.** Points represent the percentage accuracy of each tool in different expansion length bin, aggregated across all R10.4.1 samples.

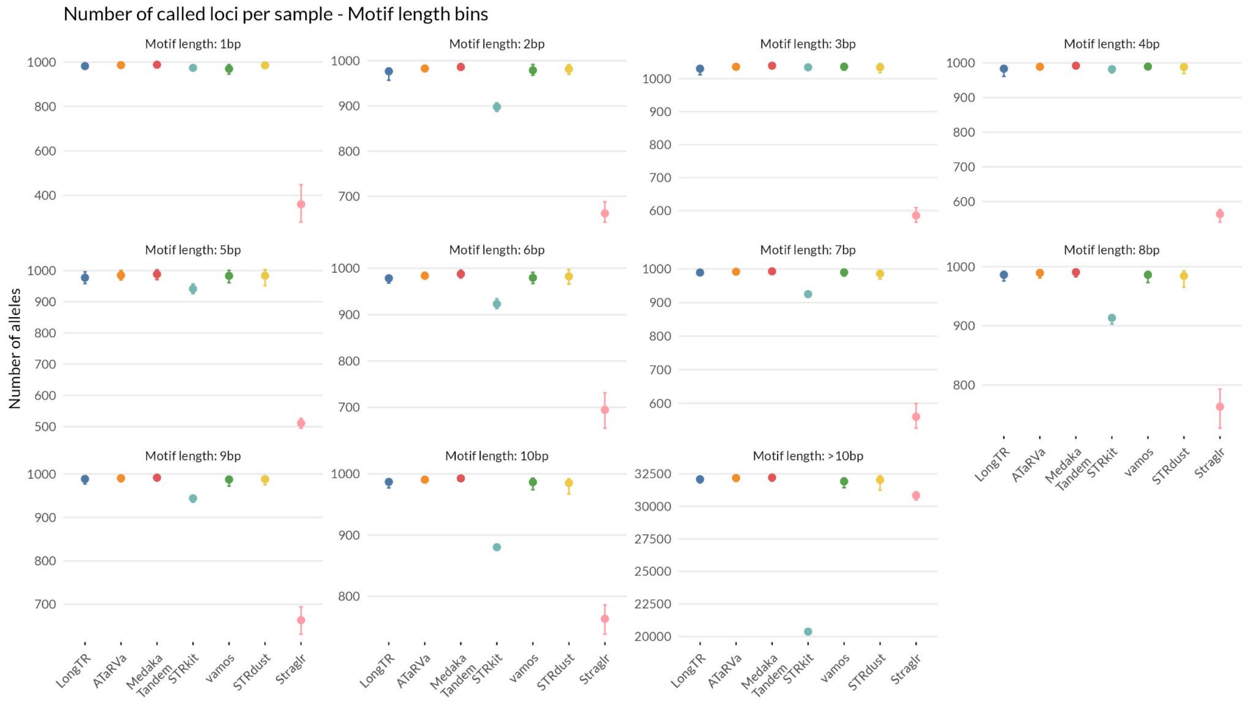


**Supplementary Fig 6**: **Average number of loci called per motif length category across all samples.** Points represent the mean number of loci called by each tool across all samples for motif length categories spanning 1–10 bp and >10 bp. Error bars indicate the range between the minimum and maximum number of calls observed across samples within each category.

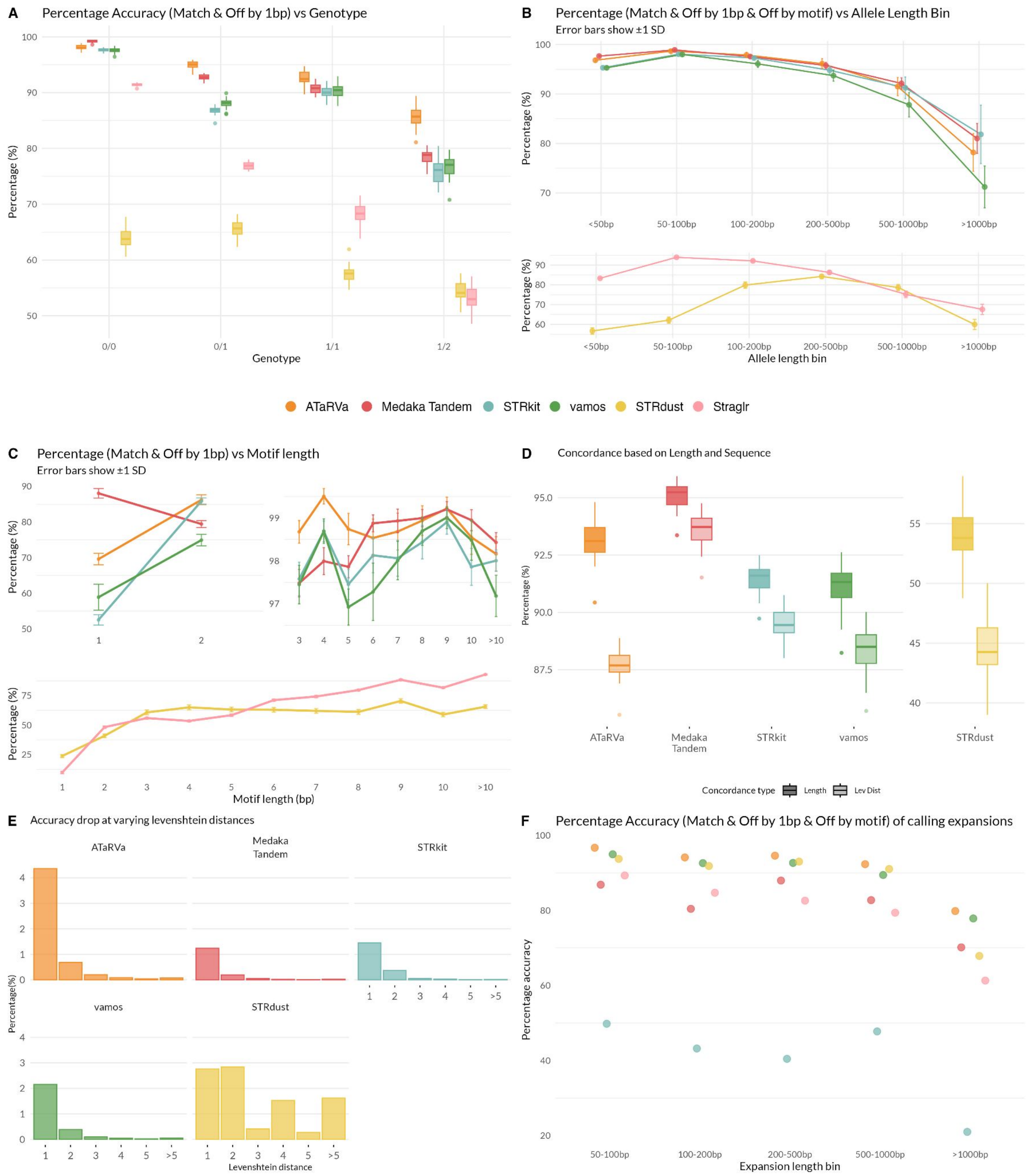


**Supplementary Fig 7**: **Accuracy of genotypers across different stratifications calculated as the percentage of allele calls with length within 1 bp with the assembly for HPRC R9.4.1 samples. A.** Accuracy of each tool stratified by genotype class: homozygous reference (0/0), heterozygous reference (0/1), homozygous alternate (1/1), and heterozygous alternate (1/2). Points represent mean accuracy per genotype class across all samples. Points represent mean accuracy, and error bars indicate ±1 standard deviation across samples. **B.** Accuracy of each tool across allele length bins (<50 bp, 50–100 bp, 100–200 bp, 200–500 bp, 500–1000 bp, and >1000 bp) in R9.4.1 samples. Points represent mean accuracy, and error bars indicate ±1 standard deviation across samples. Tools are displayed in two separate panels for readability. **C.** Accuracy of each tool across motif length categories (1–10 bp and >10 bp) in R9.4.1 samples. Points represent mean accuracy, and error bars indicate ±1 standard deviation across samples. Tools are displayed across three panels for readability. The upper two panels show the five higher-performing tools (LongTR, ATaRVa, Medaka Tandem, STRkit, and vamos), with mononucleotide and dinucleotide repeats shown separately on a y-axis range of 50–95% and motif lengths ≥3 bp displayed on an expanded y-axis of 98–100%. The lower panel shows STRdust and Straglr on a broader scale to capture their wider accuracy range across all motif lengths. **D.** Accuracy calculated for each tool on two levels: length-based accuracy, considering all calls with a matching allele length, and sequence-based accuracy, calculated using Levenshtein distance for the subset of calls with a matching allele length. The difference between the two values reflects the proportion of length-concordant calls that differ at the sequence level from the assembly allele. **E.** Contribution of Levenshtein distance categories to sequence-level accuracy drop. Bar plots showing the percentage of total calls at each Levenshtein distance value for each tool, restricted to calls with matching allele lengths. Each bar represents the proportion of calls at a given Levenshtein distance, illustrating the relative contribution of different magnitudes of sequence deviation to the overall sequence-level accuracy drop observed for each tool. **F.** Accuracy of expansion calls per tool across R9.4.1 samples. Points represent the percentage accuracy of each tool in different expansion length bin, aggregated across all R9.4.1 samples.

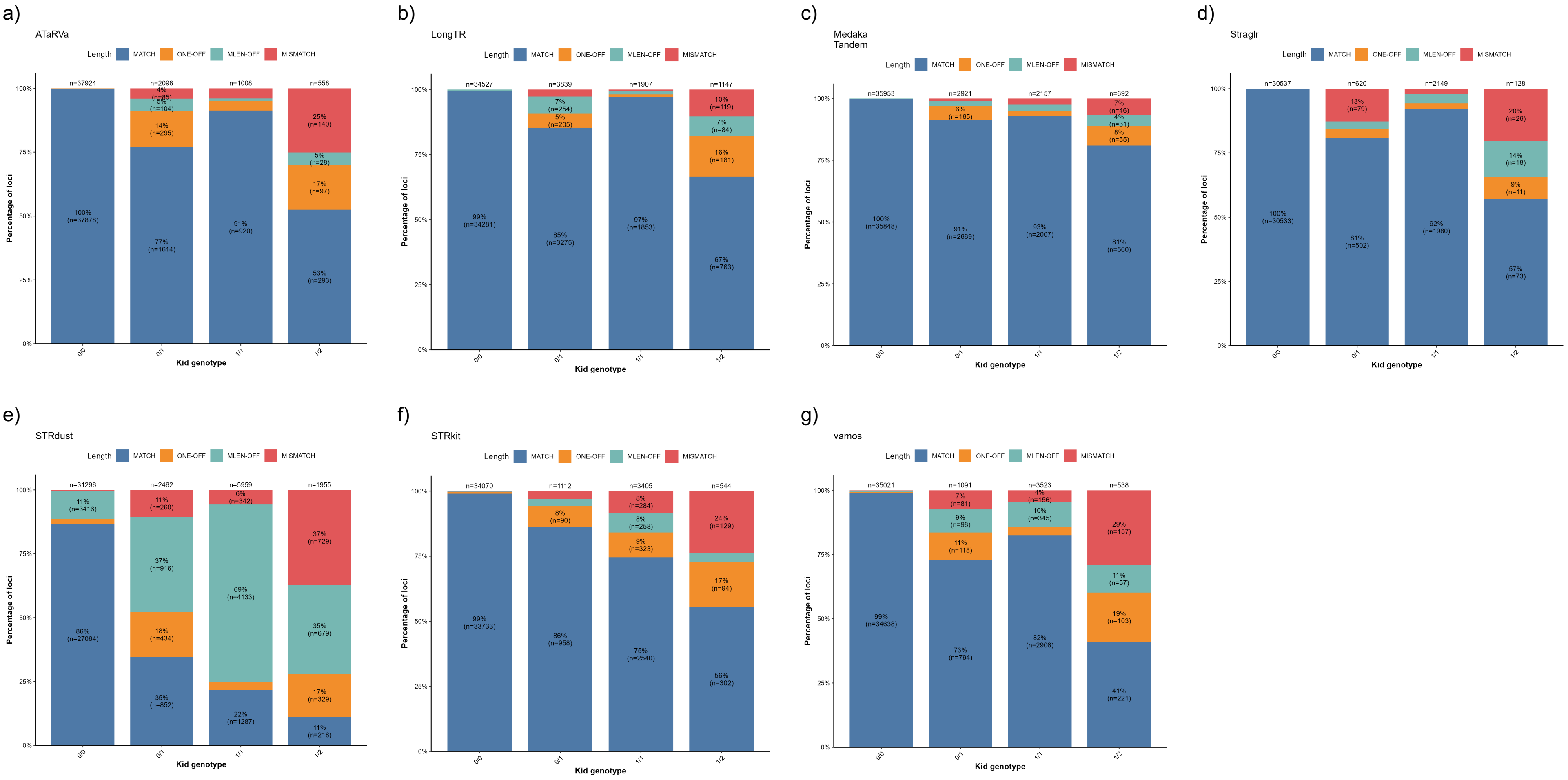


**Supplementary Fig 8. Mendelian consistency of tandem repeat genotyping tools across genotype classes in the HG002 GIAB trio.** Stacked bar charts showing the proportion of loci in each Mendelian consistency category for seven TR genotyping tools (ATaRVa, LongTR, Medaka Tandem, Straglr, STRdust, STRkit, and vamos), stratified by child genotype class (0/0, 0/1, 1/1, and 1/2) in the HG002 GIAB trio. Each bar represents all loci assigned to a given genotype class, with the total locus count shown above. Loci are classified into four categories: MATCH (child allele lengths exactly consistent with parental alleles); ONE-OFF (consistent within ±1 bp); MLEN-OFF (consistent within less than one full repeat motif unit); and MISMATCH (not consistent under any of the above criteria). Tools are arranged alphabetically.

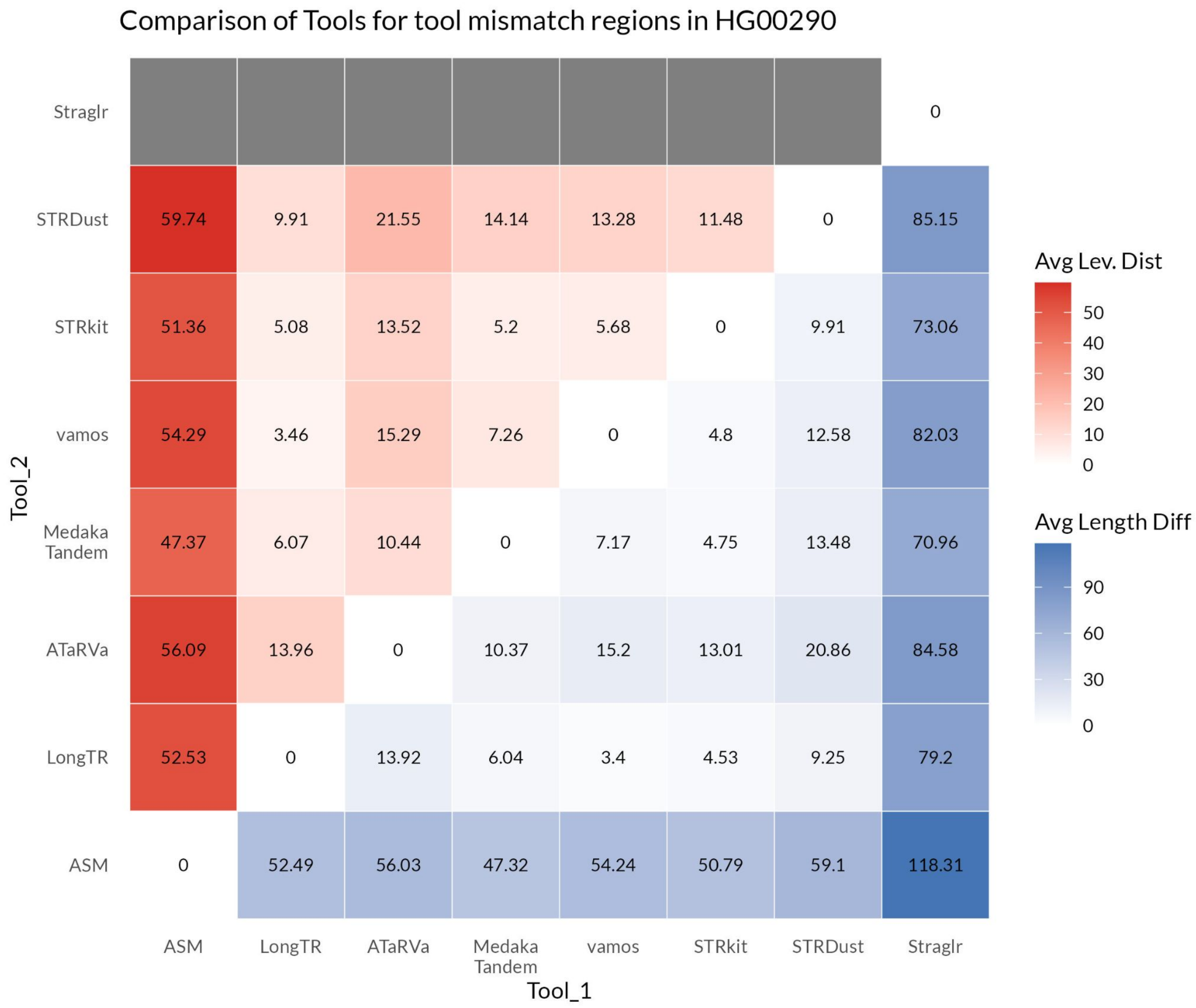


**Supplementary Fig 9**: **Pairwise comparison of genotyping tools for sample HG00290 at assembly discordant loci.** The heatmap displays pairwise comparisons between all tools, including genome assemblies as a reference genotyping method for sample HG00290, restricted to loci where at least five tools disagree with the assembly genotype. The upper triangle represents the mean Levenshtein distance between allele sequences, and the lower triangle represents the mean absolute length difference between alleles, averaged across all commonly called loci and samples. Straglr is excluded from sequence-level comparisons as it does not report allele sequences

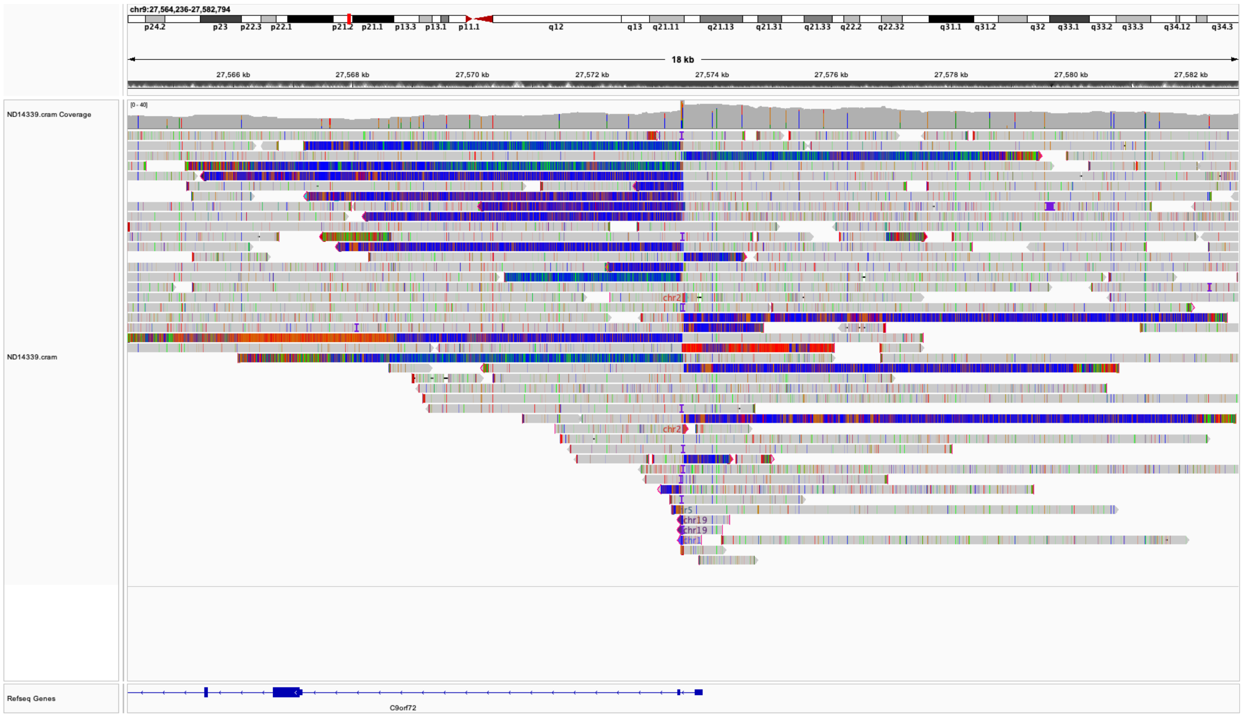


**Supplementary Fig 10:** The C9orf72 expansion in sample ND14339 was missed by all genotypers. While there was evidence of an expansion in the soft-clips, the aligner did not identify any spanning reads, and potentially spanning soft-clipped reads were low-quality.
